# Impact of innate immune activation on T cell dynamics and functional recovery following traumatic brain injury

**DOI:** 10.64898/2026.03.23.713833

**Authors:** Sahil Threja, Nathan Ryzewski Strogulski, Janeen Laabei, Gloria Vegliante, Carly Douglas, Tizbit Ashine Bogale, Ciara Moynihan, Giusy Di Franco, Matthias Mack, Lisa Borkner, Béré Diallo, Kingston H.G. Mills, David J. Loane

**Author notes:** Address correspondence to: David Loane Ph.D., School of Biochemistry & Immunology, Trinity College Dublin, Room 5.08C, 152-160 Pearse Street, Dublin, D02R590, Ireland. **Competing interest declaration:** The authors declare no competing interests. **Data availability declaration:** Data are available upon request and can be accessed through the corresponding author. **Ethics, consent to participate, and consent to publish declarations:** Not applicable.

## Abstract

**Background:** Traumatic brain injury (TBI) initiates a rapidly evolving neuroinflammatory response; however, the temporal relationship between early innate immune activation, T cell polarization, and neurobehavioural recovery remains poorly understood. Here, we hypothesize that interleukin-1β (IL-1β) is a critical upstream mediator that polarizes T cells towards pro-inflammatory and cytotoxic effector functions following TBI.

**Methods:** Using a controlled cortical impact model in adult male C57BL/6J mice, we mapped post-injury immune dynamics and investigated whether targeting key innate inflammatory compartments influenced subsequent T cell programming and neurological outcomes. We conducted longitudinal immune profiling by multiparameter spectral flow cytometry and quantitative polymerase chain reaction up to 10 days post-injury. Antibody-based immune depletion strategies were used to investigate neutrophil and monocyte contributions to the post-traumatic T cell response, while pharmacological inhibition of NLRP3 inflammasome by MCC950 treatment was used to investigate the contribution of IL-1β.

**Results:** TBI elicited a structured early innate immune response, marked by rapid chemokine induction, followed by temporally distinct infiltration of neutrophils, monocytes, and dendritic cells. Neutrophils and monocytes were the predominant early IL-1β-producing infiltrating populations. This was followed by a delayed adaptive phase characterized by sustained recruitment of T cell subsets (CD4+, CD8+, γδ+), alongside dynamic effector cytokine production (IL-17, IFN-γ). Neutrophil depletion altered the early myeloid composition but did not result in durable improvements in T cell effector responses or neurobehavioral outcomes. Depletion of CCR2-dependent inflammatory monocytes reduced acute monocyte accumulation and attenuated early downstream T cell responses; however, these effects were not sustained and only resulted in modest neurobehavioural benefits. In contrast, inhibition of the NLRP3 inflammasome suppressed microglial IL-1β production, without significantly altering leukocyte recruitment or subacute T cell effector phenotypes. These phenotypic changes were associated with improvements in motor and cognitive function recovery.

**Conclusion:** We show that early monocyte IL-1β signalling actively regulates downstream T cell infiltration and effector function after TBI. In addition, inhibition of NLRP3 inflammasome after TBI attenuates microglial IL-1β-associated immune activation and results in behavioural improvement despite ongoing leukocyte recruitment, indicating that targeting the nature and cellular source of IL-1β signalling can dissociate immune cell burden from neurological outcomes. Collectively, our findings identify myeloid IL-1β-linked pathways as a viable bridge between innate and adaptive immunity post-TBI, and underscore cellular compensation as a critical design consideration for next-generation immunotherapies.

## Introduction

Traumatic brain injury (TBI) is a major cause of mortality and prolonged neurological disability worldwide [1], largely due to the progression of secondary injury cascades that evolve over time ranging from hours to months to years following the initial biomechanical brain impact. A significant contributor to this secondary pathology is neuroinflammation, which is triggered by damage-associated molecular patterns (DAMPs) that activate brain resident microglia and rapidly engage the peripheral immune system through chemokine production, blood-brain barrier (BBB) dysfunction, and leukocyte recruitment [2]. The post-traumatic neuroimmune response is not a singular event but a coordinated, time-dependent molecular and cellular program that can facilitate debris clearance and tissue repair [2]. However, when neuroinflammation is not resolved and becomes prolonged, it can exacerbate synaptic dysfunction, contribute to chronic neurodegeneration, and can be responsible for long-term poor neurological recovery in TBI patients [2]. It may even lead to the development of Alzheimer’s Disease and Related Dementias (ADRD) [3]. Interleukin-1 (IL-1) family signalling and inflammasomes are upstream amplifiers of post-traumatic neuroinflammation [4]. Specifically, the NLRP3 inflammasome regulates the maturation of IL-1β and related inflammatory signalling molecules across experimental TBI models [5-8], and has been proposed as both a biomarker and a viable therapeutic target for TBI [9]. Translational research has increasingly concentrated on IL-1 pathway modulation as a therapeutic strategy to mitigate harmful neuroinflammation without completely suppressing host defence, with growing research efforts to optimize timing, compartment, and TBI patient stratification to the therapeutic window [4].

However, accumulating evidence indicates that adaptive immunity is not merely a late bystander in secondary neuroinflammation following TBI. Complex T and B cell responses can persist into subacute and chronic phases and can influence myeloid phenotypes and neurological outcomes [10]. The manner in which early, myeloid-dominated inflammation instructs subsequent T cell effector programs remains incompletely understood, particularly when multiple innate compartments may compensate for one another. In this study, we used a clinically relevant model of experimental TBI (i.e. controlled cortical impact, CCI) in adult male wildtype C57BL/6J mice to map the kinetics of innate leukocyte recruitment and IL-1β production, and subsequently investigated how perturbing key upstream cellular and molecular nodes, including neutrophils, CCR2-dependent monocytes, and NLRP3 inflammasome signalling, reshapes the evolving adaptive immune landscape and influences neurological outcomes in TBI mice.

## Methods

### Animals

Adult male C57BL/6J mice (10-12 weeks old; 24-27g) were bred in-house by the Comparative Medicine Unit, Trinity College Dublin, and housed in groups of five per cage under specific pathogen-free conditions (12h light/dark cycle, 24±1°C, 55±5% humidity) with *ad libitum* access to food and water. All procedures were conducted in compliance with EU regulations under oversight of the Health Products Regulatory Authority (HPRA), Ireland, designed in accordance with ARRIVE guidelines, and approved by the Trinity College Dublin Animal Research Ethics Committee and HPRA (Protocol Number: AE19136/P138).

### Controlled cortical impact

CCI was performed using a precise impactor device (RWD; Cat# 6809II) as previously described [11]. Anaesthesia was induced with isoflurane (3% induction, 0.75-1.5% maintenance), and bupivacaine (0.083%) was administered subcutaneously near the incision site. Mice were placed in a stereotaxic frame, the surgical site was cleansed with ethanol and chlorhexidine, and a 10 mm midline incision was made over the skull with the skin and fascia reflected. A 5 mm craniotomy was performed on the central aspect of the left parietal bone under surgical anaesthesia and sterile conditions. Moderate-level TBI was induced using a 3mm bevelled impactor tip at 3 m/s velocity, 1.5 mm deformation depth, and 0.18 m/s dwell time. Immediately following CCI, bupivacaine (0.083%) was administered at the injury site, the incision was closed with interrupted 5−0 silk sutures, and saline (1ml) was administered subcutaneously on the dorsal side of the mouse. Sham animals underwent the same procedure as injured mice except for craniotomy and cortical impact.

### Antibody and drug administration

A double antibody-based strategy was used for neutrophil depletion [12]. Mice were administered intraperitoneal (i.p.) injections of anti (α)-Ly6G (clone 1A8; 100 μg/mouse; Assay Genie, Cat# IVMB0025) in conjunction with a secondary antibody anti (α)-rat κ immunoglobulin light chain (clone MAR18.5; 100 μg/mouse; Assay Genie, Cat# IVMB0305). For endpoints at 3 days post-injury (dpi), antibodies were administered on −1, 1 and 2 dpi. For endpoints at 10 dpi, an additional dose was administered on day 3. Control animals received matched isotype controls for α-Ly6G and α-rat κ immunoglobulin light chain (mouse IgG1, κ and rat IgG2a, κ; Assay Genie, Ca# IVMB0198). For the depletion of CCR2+ monocytes, α-CCR2 monoclonal antibody (MC-21, 50 μg/mouse, [13]) was administered i.p. on day -1 and 1 dpi for the 3 dpi endpoint studies, with an additional dose at 3 dpi for the 10 dpi endpoint studies. Control animals received matched isotype rat IgG2b (Assay Genie, #IVMB0200) antibody. For NLRP3 inflammasome inhibition, MCC950 (10 mg/kg; MedChemExpress, Cat# HY-12815A) was administered i.p. beginning 2h post-injury, followed by repeat doses at 12, 24, and 48h post-injury for the 3 dpi endpoint studies. Dosing was continued every other day beyond 48h for the 10 dpi endpoint studies. Control animals received Vehicle (saline; i.p.).

### Neurobehavioural testing

Prior to experiments, mice were handled for 5 min per day over a minimum 5 day period. Before testing, mice were acclimatized in the behaviour room for 30-60 min. The behaviour room was maintained with uniform illumination at 35 lx, as verified by a luminometer (Extech LT300 Light Meter), and a constant noise level of 45 dB generated by a white noise machine and confirmed by a decibel meter (Extech Digital Sound Level Meter, Cat# 407730). Neurobehavioral assessments were video-tracked using ANY-maze cameras and software (Stoelting Europe).

### Beamwalk

Fine motor coordination was assessed using a beamwalk test as previously described [14]. Briefly, mice were positioned at one end of a wooden beam (5 mm width, 1300 mm length; elevated 40 cm above foam padding), and the number of foot faults made by the right hindlimb was recorded over 50 steps as the mouse progressed towards an escape house at the opposite end of the elevated beam. The mice underwent training for three consecutive days prior to sham/CCI. Mice with more than 10 foot faults at day 0 did not satisfy the basic beamwalk performance criteria and were excluded. To summarize beamwalk performance throughout the recovery period, area under the curve (AUC) was computed for each animal from day 1 onwards using the trapezoidal rule.

### Rotarod

Gross motor function was tested by an accelerating rotarod as previously described [15]. Two protocols were used. A) Standard accelerating rotarod where the rod accelerated from 4-to-60 rpm over 180 s (achieving maximum speed at 180 s). Mice underwent training for three consecutive days prior to sham/CCI, followed by daily testing post-injury (up to 3 dpi). B) Forced accelerating rotarod where the rod accelerated from 4-to-60 rpm over 20 s (achieving maximum speed at 20 s). This was a once off test with no training. During each trial, mice were placed on the rotating rod, and the latency to fall was recorded. Trials were concluded if a mouse fell, or clung to the rod and passively rotated for two complete revolutions. Each mouse completed three trials per day, with a 10 min inter-trial interval during which they had access to food and water. The mean of the three trials was used as the daily score for each mouse.

### Two-trial Y-maze

To assess spatial working memory, the two-trial Y-maze test was conducted as previously described [16] . During trial 1 (acquisition), one arm was randomly selected and blocked using a guillotine door, while the mouse was placed in a randomly chosen start arm and permitted to explore for 5 min. The mice were returned to a holding cage for a 1h inter-trial interval. In trial 2 (test), the guillotine door was removed, granting access to all three arms, and the mouse was reintroduced to the same start arm for another 5 min exploration period. Novel arm preference was calculated as follows: time in arm (new or old) / (time in new + time in old). A significant preference of the new over the old arm was considered indicative of intact working memory.

### The Simple Neuroassessment of Asymmetric imPairment (SNAP)

Testing was conducted in mice as previously described [17], assessing eight distinct parameters: handler interaction, grip strength, visual placing, pacing or circling, gait and posture, head tilt, visual field, and coordination and proprioception. Each parameter was rated on a scale from 0 (normal) to 5 (severely impaired), and the cumulative score, ranging from 0 (best) to 40 (worst), was derived by summing these individual scores.

### RT-Qpcr

Total RNA was extracted from snap-frozen sham and TBI ipsilateral hippocampal tissue punches as previously described [18]. Total RNA was isolated using a RNeasy Mini Kit (Qiagen; Cat# 74104) and cDNA was prepared using the Verso cDNA Synthesis Kit (ThermoFisher Scientific; Cat# 4368814). Real-time polymerase chain reaction (RT-qPCR) amplification was conducted using the 7500 Fast Real-Time PCR System with SYBR Green PCR Master Mix (CliniScience, Cat# Q711-03) and the corresponding primers. The specific primer sequences were as follows: CXCL1: forward 5′-CACCCAAACCGAAGTCATAGCC-3′, reverse 5′-AAGCCAGCGTTCACCAGACA-3′; CCL2: forward 5′-GTTGGCTCAGCCAGATGC-3′, reverse 5′-AGCCTACTCATTGGGATCATCTTG-3′; CXCL10: forward 5′-TTTCTGCCTCATCCTGCT-3′, reverse 5′-TCCCTATGGCCCTCATTCT-3′; IL-1b: forward 5′-GAGGACATGAGCACCTTCTTT-3′,reverse 5′-GCCTGTAGTGCAGTTGTCTAA-3′; ICAM-1: forward 5′-AAACCAGACCCTGGAACTGCAC-3′, reverse 5′-GCCTGGCATTTCAGAGTCTGCT-3′; ITGB2: forward 5′-CTTTCCGAGAGCAACATCCAGC-3′, reverse 5′-GTTGCTGGAGTCGTCAGACAGT-3′; GAPDH: forward 5′-TGGTGAAGGTCGGTGTGAAC-3′, reverse 5′-TGAATTTGCCGTGAGTGGAG-3′; β-actin: forward 5’-GCC TTC CTT CTT GGG TAT GG -3’, reverse 5’-AGC ACT GTG TTG GCA TAG AG -3’. Gene expression levels were normalized to the endogenous control β-actin/GAPDH mRNA level and are presented as fold change relative to the sham group.

### Flow cytometry

Mice were humanely euthanized via isoflurane overdose and were perfused through the left ventricle with 30ml of cold PBS prior to brain extraction. The ipsilateral cortex, excluding the olfactory bulb and cerebellum, was sectioned and collected into 1ml of cRPMI. DNase (Merck, Cat# D4263-1VL) and Collagenase (Merck, Cat# 11088866001) (10µg/ml) were added to the tubes and incubated at 37°C for 30 min on a shaker (Incu-Shaker mini, Benchmark Scientific). The homogenate was passed through a 70µm cell strainer, supplemented with cRPMI, centrifuged, and resuspended in 40% Percoll before being layered over 70% Percoll and centrifuged with brakes off (600 × g; 20min, 4°C). The upper myelin layer was discarded, and isolated mononuclear cells were collected from the interface of the Percoll gradient. Cells were washed with cRPMI and plated on a 96-well V-plate, then immediately stained for flow cytometry. Following incubation with Brefeldin A (5µg/ml, Sigma Aldrich; Cat #B7651), PMA (20ng/ml, Merck; Cat #P1585) and Ionomycin (0.5µg/ml, Merck; Cat # 554724) for 3h at 37°C, cells were centrifuged and stained with Live/Dead (Invitrogen; 50µl/well diluted 1:600; Cat# L34957; 10min; dark, room temperature). Cells were washed with FACS buffer and resuspended in a master mix of mouse Fc Block (BD Pharmingen; Cat# 553141) and surface fluor-conjugated antibodies (1:200–400 dilutions): CD19-BV570 (Cat# 115535), Ly6C-BV605 (Cat # 563011), Ly6G-BV650 (Cat # BL-127641), CD45-BV750 (Cat# 103157), MHC-II (Cat# 107645), CD3e-BB700 (Cat# 566494), TCRγδ-PerCP-ef710 (Cat # 46-5711-82), CD4-PeCy-5 (Cat #15-0041-82), CD8-APC (Cat # 17-0081-82), CD11b-APCeF780 (Cat # 47-0112-82), and NK1.1-AF700 (Cat # 56-5941-82). Following a wash step, cells underwent intracellular staining by adding Fixation/Permeabilization working solution and incubated for 20min at room temperature. Cells were washed twice with 1X Permeabilization buffer and centrifuged (1350 RPM; 5min, 4°C). The pelleted cells were resuspended in the intracellular antibody master mix at a concentration of 1:200, containing IL-17A-Pacific blue (Cat# 506918), Granzyme-B-FITC(Cat# 11-8898-82), IFN-γ-PE-CF594 (Cat# 562303), and IL-1β-PE-Cy7 (Cat# 25-7114-80) in 1X Permeabilization buffer and incubated overnight at 4°C (in dark). Cells were washed twice in 1X Permeabilization buffer and finally washed in FACS buffer for analysis. All flow cytometric data were acquired using Cytek Aurora Spectral flow cytometry with SpectroFlo software. Compensation was calculated using single-stained cells. Cell-specific fluorescence minus one (FMO) controls were employed for gating strategies, and flow cytometric analysis was performed using FloJo software (TreeStar Inc.). UMAP (Uniform Manifold Approximation and Projection) was applied to concatenated, down sampled live singlet events using the FlowJo UMAP plugin and a marker set restricted to phenotypic channels of interest (excluding FSC/SSC, time, and viability channels). UMAP settings (e.g., number of nearest neighbours and minimum distance) were held constant across samples within an analysis to enable direct comparison of embeddings. Cell populations were identified using the gating strategy shown in **Suppl. Fig.1**. The absolute number of cells was calculated based on the frequency of the total parent population.

### Rigor, reproducibility, and statistical analyses

Mice were randomly allocated to groups using a computer-generated sequence (excel). The experimental unit was the individual mouse. Inclusion and exclusion criteria were defined a priori, and any exclusions were reported with justifications. Sample sizes were determined based on prior data and/or an a priori power calculation focused on the primary endpoint. The individual responsible for drug administration was blinded to treatment allocation, and behavioural analyses were performed by investigators blinded to both injury and treatment. Statistical analysis was performed using Prism v9.1.0 software (GraphPad Software, San Diego, CA, USA; RRID: SCR_002798). Normality testing was conducted using the Shapiro-Wilks test, and parametric statistical analysis was applied where data conformed to normality. Data were analysed using paired and unpaired Student’s t-tests, one-way ANOVA with post hoc Dunnett’s multiple comparisons test, or two-way ANOVA, with or without repeated measures, employing post-hoc Tukey’s test to correct for multiple comparisons. All quantitative data are presented as mean ± standard error of the mean (SEM). p-values less than 0.05 were considered statistically significant.

## Results

### TBI produces a rapid interleukin-1β production in infiltrating myeloid cells

To investigate the temporal activation of infiltrating innate immune cells following TBI, we performed a longitudinal analysis of immune gene expression and leukocyte composition within the injured hemisphere. Mice were subjected to either moderate-level CCI or sham surgery and hippocampal tissue punches were collected at 3, 6, 12, 24 hours post-injury (hpi) for quantitative PCR; the remaining ipsilateral tissues (excluding the olfactory bulb and the cerebellum) was used for multiparameter flow cytometry analysis of infiltrating innate immune cells (**Fig. 1A**). qPCR analysis revealed a rapid induction of chemokine *Cxcl1* expression in the ipsilateral hippocampus, which peaked at 3 hpi compared to sham (**Fig. 1B**), while *Ccl2* gene induction was delayed with high expression at 24 hpi (**Fig. 1C**). The profile of hippocampal *Il1b* gene expression mirrored Cxcl1 and peaked at 6 hpi (**Fig. 1D**). In the same injured tissues, the numbers of IL-1 β ^+^ infiltrating cells were quantified. Representative flow cytometry plots for the infiltrating innate immune cells are depicted in (**Fig. 1E**), and a back-gating strategy was used to quantify IL-1 β production in each subset. There was a pronounced early infiltration of Ly6G^+^ neutrophils at 6 hpi compared to sham (**Fig. 1E,G**), whereas Ly6C^+^ monocytes and CD11c^+^ dendritic cells peaked at 12 hpi (**Fig. 1E,H,I**). Neutrophils and monocytes were the major IL-1 β -producing cells (**Fig.1F,J,K**). In addition, there was the expected increase in brain resident microglial proliferation following TBI (**Suppl. Fig 2**). UMAP analysis of innate immune cell populations and cytokine production throughout this early post-injury time course (**Suppl. Fig. 3**) demonstrate that TBI elicits a rapid and structured innate immune cascade, with early transcriptional chemokine activation giving rise to temporally distinct recruitment of neutrophils and monocyte-lineage cells that contribute to IL-1β production in the acute post-injury period.

**Figure 1.**
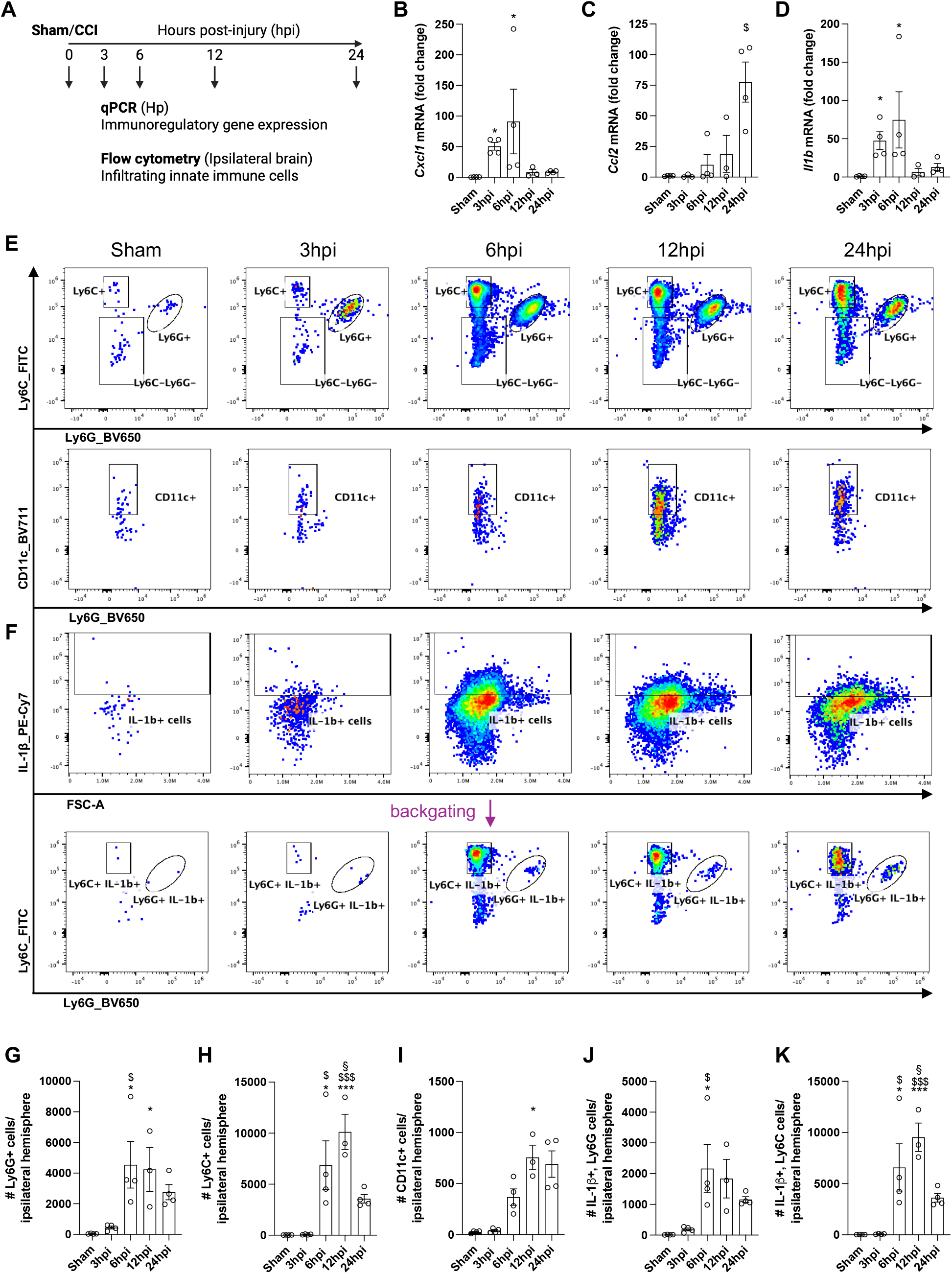
Early infiltration of IL-1β-producing innate immune cells into the injured brain. (A) Experimental design. (B-D) Relative fold-change hippocampal gene expression of *Cxcl1, Ccl2, Il1b*. (E) Representative flow cytometry plots for Ly6C^+^ monocytes, Ly6G^+^ neutrophils and CD11c^+^ dendritic cells. (F) Back-gating strategy to identify IL-1β-producing innate immune cells. (G-K) Bar graphs illustrate the absolute numbers of Ly6G^+^ neutrophils (G), Ly6C^+^ monocytes (H), CD11c^+^ dendritic cells (I), IL-1β-producing neutrophils (J), IL-1β-producing monocytes (K). Data = Mean ± SEM (n=3-4 per time point); differences vs. Sham *p<0.05, **p<0.01, ***p<0.001; differences vs. 3hpi ^$^p<0.05, ^$$^p<0.01, ^$$$^p<0.001; differences vs. 6hpi ^*#*^p<0.05, ^##^p<0.01, ^###^p<0.001; differences vs. 12hpi ^§^p<0.05, ^§§^p<0.01, ^§§§^p<0.001 by one-way ANOVA with Tukey’s multiple comparisons test. Abbreviations: Hp, hippocampus.

### TBI drives pro-inflammatory lymphocyte recruitment during the subacute phase post-injury

Following the initial innate immune cell infiltration, adaptive immune cells traffic to injured brain parenchyma in response to chemokine gradients and a damaged BBB. We conducted a more extensive and chronic time course to quantify immunoregulatory gene expression and T cells at 1, 3, 10, and 28 days post-injury (dpi) using the prior experimental approaches (**Fig. 2A**). There was a significant elevation in *Cxcl10, Icam1* and *Itgb2* mRNA, which peaked at 3 dpi, remained elevated until 10 dpi, and returned to baseline by 28 dpi compared to sham (**Fig. 2B-D**). In the same injured tissues, the numbers of IL-17^+^ and IFN-γ^+^ infiltrating lymphocytes were quantified. Representative flow cytometry plots for the infiltrating adaptive immune cells are presented (**Fig. 2E**), while representative plots for cytokine production from these subpopulations are shown (**Fig. 2F**). Flow cytometric quantification revealed peak infiltration of CD4^+^ (**Fig. 2G**), CD8^+^ (**Fig. 2H**), and NK^+^ (**Fig. 2I**) T cells at 10 dpi compared to sham, while γδ^+^ (**Fig. 2J**) T cells peaked earlier at 3 dpi. The production levels of IFN-γ mirrored the infiltration trends for CD4^+^ (**Fig. 2K**), CD8^+^ (**Fig. 2L**), and NK^+^ (**Fig. 2M**) T cells. However, γδ^+^ T cells were the exclusive producers of IL-17 (**Fig. 2N**), with numbers increasing at 1 dpi and significantly upregulated at 3 dpi compared to sham. In contrast, IFN-γ production by γδ^+^ T cells (**Fig. 2O**) peaked at 10 dpi. There was the expected increase in brain resident microglial proliferation following TBI and numbers remained elevated through 28 dpi (**Suppl. Fig 2**). UMAP analysis of adaptive immune cell populations and cytokine production throughout this chronic post-injury time course (**Suppl. Fig. 4**) demonstrate a temporally coordinated adaptive immune response to TBI, characterized by sustained chemokine and adhesion molecule induction, progressive accumulation of T cell subsets, and distinct pro-inflammatory cytokine production profiles that peak during the subacute phase post-injury.

**Figure 2.**
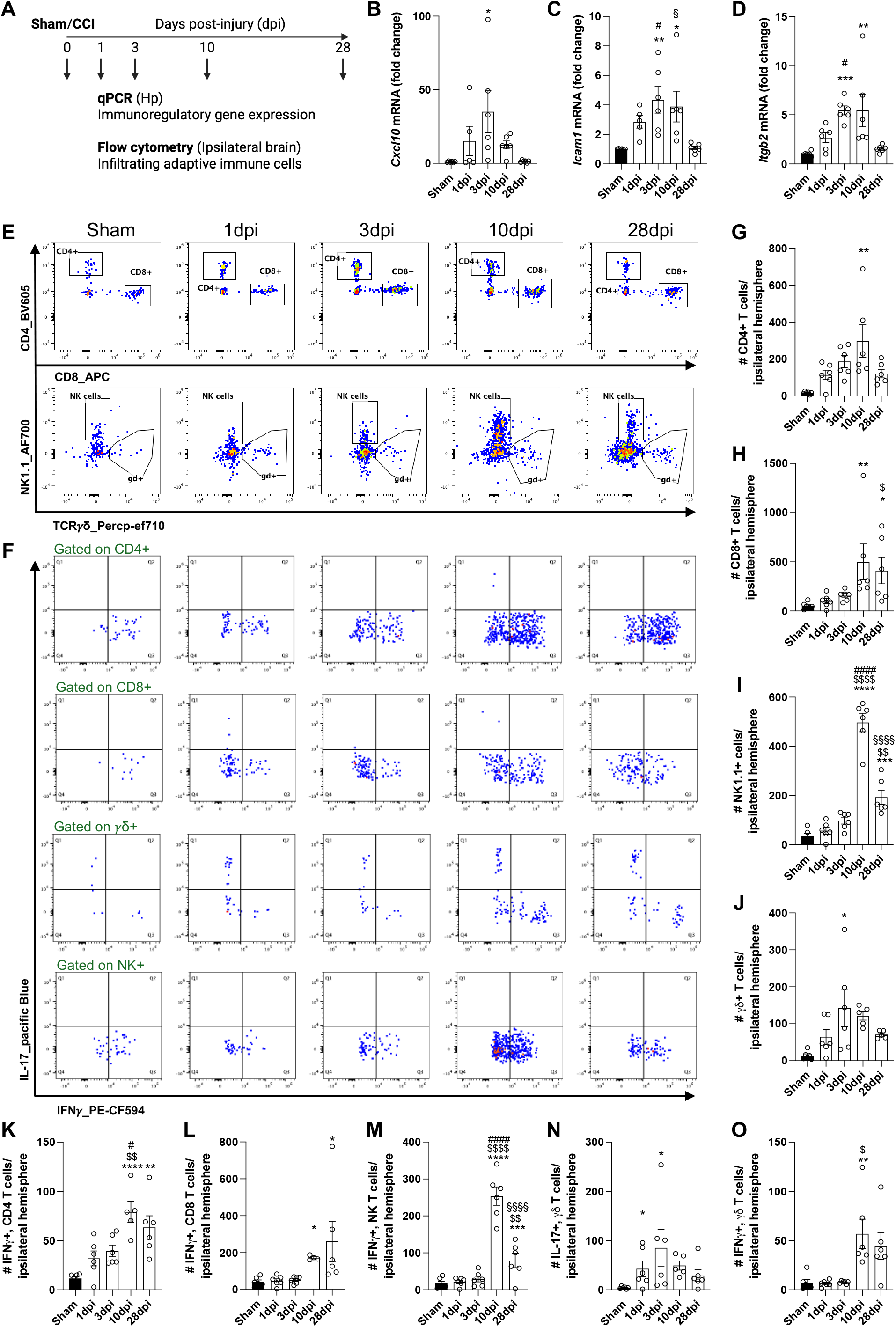
Sustained infiltration of T cells and pro-inflammatory cytokine production in the subacute phase post-injury. (A) Experimental design. (B-D) Relative fold-change expression levels of *Cxcl10, Icam1, Itgb2*. (E) Flow cytometry plots of CD4^+^, CD8^+^, γδ^+^, and NK^+^ T cells. (F) Representative flow cytometry plots for IL-17 and IFN-γ production by different T cell subsets. (G-O) Bar graphs illustrate the absolute numbers of CD4+ T cells (G), CD8+ T cells (H), NK^+^ T cells (I), γδ^+^ T cells (J), IFN-γ-producing CD4^+^ T cells (K), IFN-γ-producing CD8^+^ T cells (L), IFN-γ-producing NK^+^ T cells (M), IL-17-producing γδ^+^ T cells (N), and IFN-γ-producing γδ^+^ T cells (O). Data = Mean ± SEM (n=6 per time point); differences vs. Sham *p<0.05, **p<0.01, ***p<0.001; differences vs. 1dpi ^$^p<0.05, ^$$^p<0.01, ^$$$^p<0.001; differences vs. 3dpi ^*#*^p<0.05, ^##^p<0.01, ^###^p<0.001; differences vs. 10dpi ^§^p<0.05, ^§§^p<0.01, ^§§§^p<0.001 by one-way ANOVA with Tukey’s multiple comparisons. Abbreviations: Hp, hippocampus.

### Neutrophil depletion increases IL-1β^+^ myeloid cell accumulation and the expansion of IL-17^+^ γδ T cells in the injured brain

Given the rapid recruitment of IL-1β^+^ neutrophils, we next investigated their role in T cell responses during the subacute phase post-injury. In proof-of-concept pilot studies we depleted neutrophils prior to TBI using a piggy-back approach [12], where anti(α)-Ly6G combined with anti(α)-rat kappa antibody was intraperitoneally (I.P.) administered to CCI mice 12h prior to TBI. Blood and brain were collected at 12h post-injury. This method resulted in a complete depletion of neutrophils in the circulation and injured brain (**Suppl. Fig. 5**). We then performed neutrophil depletion in sham and CCI mice, where the α-Ly6G/α-rat kappa antibody was administered I.P. at -1, 1 and 2 dpi. Rotarod assessment of gross motor function was performed prior to, and after TBI, and brain tissue was collected at 3 dpi for flow cytometry analysis (**Fig. 3A**).

**Figure 3.**
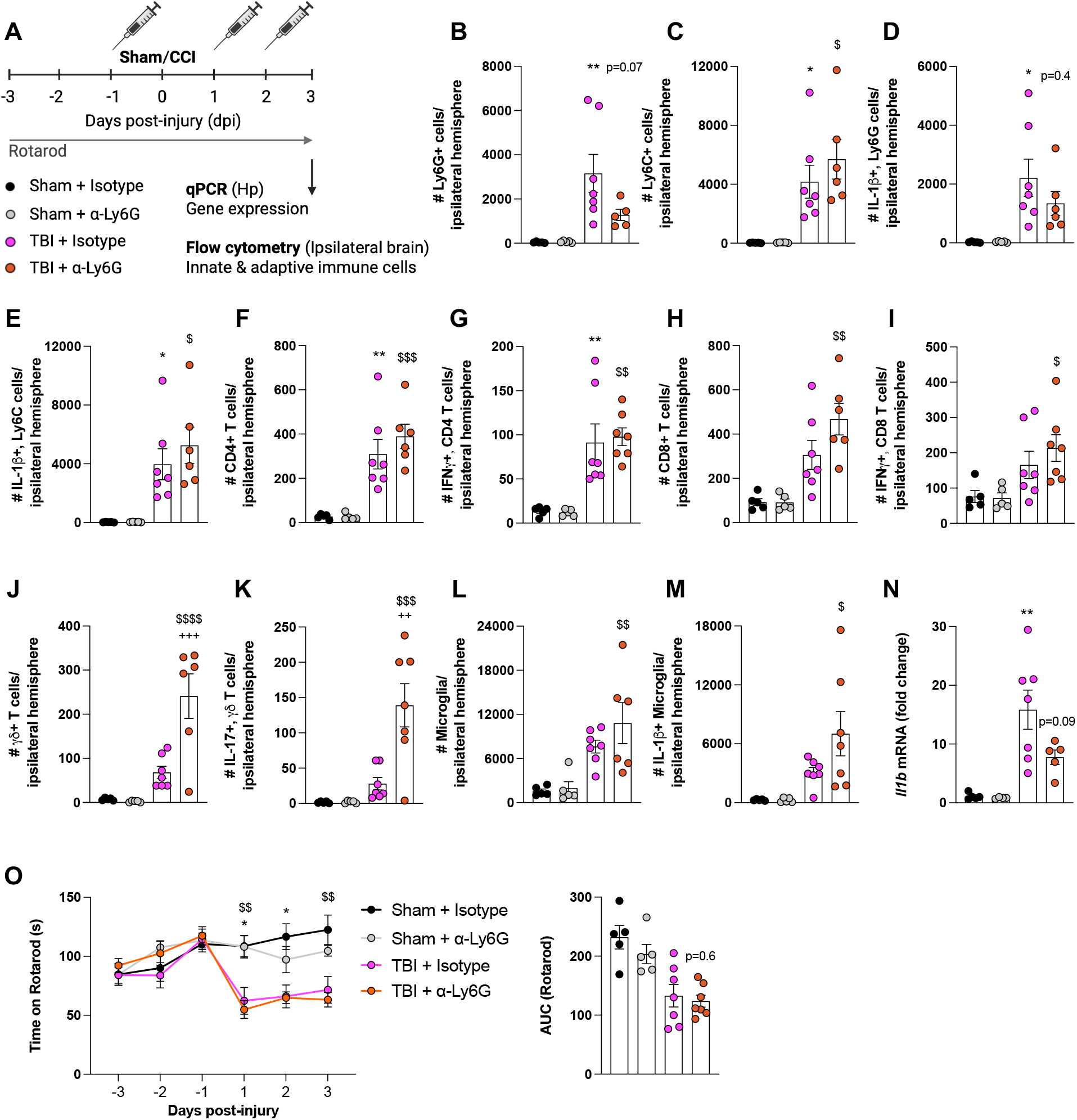
Neutrophil depletion does not alter T cell numbers and cytokine production post-TBI. (A) Experimental design. (B-M) Bar graphs illustrate the absolute numbers of Ly6G^+^ neutrophils (B), Ly6C^+^ monocytes (C), IL-1β-producing neutrophils (D), IL-1β-producing monocytes (E), CD4^+^ T cells (F), IFN-γ-producing CD4^+^ T cells (G), CD8^+^ T cells (H), IFN-γ-producing CD8^+^ T cells (I), γδ^+^ T cells (J), IL-17-producing γδ^+^ T cells (K), microglia (L), and IL-1β-producing microglia (M). Relative fold-change expression levels of *Il1b* mRNA in hippocampus (N). Neurobehavioural tests: Rotarod (O). Data presented as Mean ± SEM (n=5-7 per group); Sham + Isotype vs. TBI + Isotype *p<0.05, **p<0.01, ***p<0.001; Sham + α-Ly6G vs. TBI + α-Ly6G ^$^p<0.05, ^$$^p<0.01, ^$$$^p<0.001; TBI + Isotype vs. TBI + α-Ly6G ^+^p<0.05, ^++^p<0.01, ^+++^p<0.001 by two-way ANOVA with post-hoc Tukey’s test to correct for multiple comparisons . Abbreviations: Hp, hippocampus.

TBI resulted in a marked increase in neutrophils compared to sham, while α-Ly6G treatment led to a non-significant reduction in neutrophil numbers (p=0.07; **Fig. 3B**), indicating a partial neutrophil depletion at 3 dpi. Interestingly, the number of Ly6C^+^ monocytes was increased after TBI (**Fig. 3C**), and were even higher in the α-Ly6G treatment group. Consistent with an inflammatory phenotype, IL-1β^+^ positive neutrophils were increased post-injury (**Fig. 3D**); however, this increase was not altered by α-Ly6G treatment at this time point (p=0.4 vs. TBI + Isotype). In contrast, IL-1β^+^ monocytes were increased with TBI (**Fig. 3E**), and were further augmented by α-Ly6G treatment. The increase in CD4^+^ T cells (**Fig. 3F**) and IFN-γ^+^ CD4^+^ T cells (**Fig. 3G**), were further enhanced by α-Ly6G treatment. A similar trend was observed for CD8^+^ T cells (**Fig. 3H**) and IFN-γ^+^ CD8^+^ T cells (**Fig.3I**). Notably, the most significant change in T cell subtypes associated with α-Ly6G treatment was the expansion of γδ^+^ T cells (**Fig. 3J**) and IL-17^+^ γδ^+^ T cells (**Fig. 3K**) in TBI mice. Activation of microglia was also evident, with proliferation of microglia (**Fig. 3L**) and increased numbers of IL-1β-producing microglia (**Fig. 3M**) after TBI, and even higher numbers of IL-1β^+^ microglia with α-Ly6G treatment. *Il1b* mRNA expression in the hippocampus was increased by TBI, and this was partially reduced by α-Ly6G treatment (**Fig. 3N**). Despite these changes in infiltrating and resident immune cells in the brain, α-Ly6G treated TBI mice had similar motor performance compared to TBI mice treated with an isotype control antibody (**Fig. 3O**). Consistent with this, area-under-the-curve analysis revealed no significant treatment effect, with no difference between TBI + α-Ly6G and TBI + isotype groups (p=0.6). Collectively, these findings suggest that neutrophil depletion by α-Ly6G treatment did not provide an early functional advantage following TBI. Despite achieving only a partial reduction in Ly6G^+^ neutrophils at this time point, the α-Ly6G treatment resulted in a shift towards an enhanced inflammatory profile in the injured brain. This profile was characterized by increased IL-1β^+^ monocytes/microglia and the expansion of IL-17^+^ γδ^+^ T cells in the injured brain.

### Neutrophil depletion does not alter T cell effector phenotypes or neurobehavioral outcomes at 10dpi

We next investigated whether α-Ly6G-mediated neutrophil depletion altered T cells and outcomes up to 10 dpi (**Fig. 4A**). TBI mice exhibited a significant accumulation of CD4^+^ T cells in the injured brain at 10 dpi (**Fig. 4B**), and there was no reduction in CD4^+^ T cells with α-Ly6G treatment. IFN-γ^+^- (**Fig.4C**) and Granzyme B^+^ (GzmB)- (**Fig. 4D**) CD4^+^ T cells were also increased at 10 dpi, with comparable levels in isotype control and α-Ly6G treated TBI mice. There was a similar pattern for total CD8^+^ T cells (**Fig. 4E**), IFN-γ^+^- (**Fig. 4F**) and GzmB^+^- (**Fig. 4G**) CD8^+^ T cells; all of which were elevated post-injury regardless of neutrophil neutralization. Notably, the γδ^+^ T cell compartment remained significantly altered at 10 dpi. γδ^+^ T cells were increased post-injury (**Fig. 4H**) and were further elevated with α-Ly6G treatment, accompanied by a marked increase in IL-17^+^ -γδ^+^ T cells (**Fig. 4I**). In contrast, IFN-γ^+^-γδ^+^ T cells were present at very low levels (**Fig. 4J**), while there was no effect of α-Ly6G treatment.

**Figure 4.**
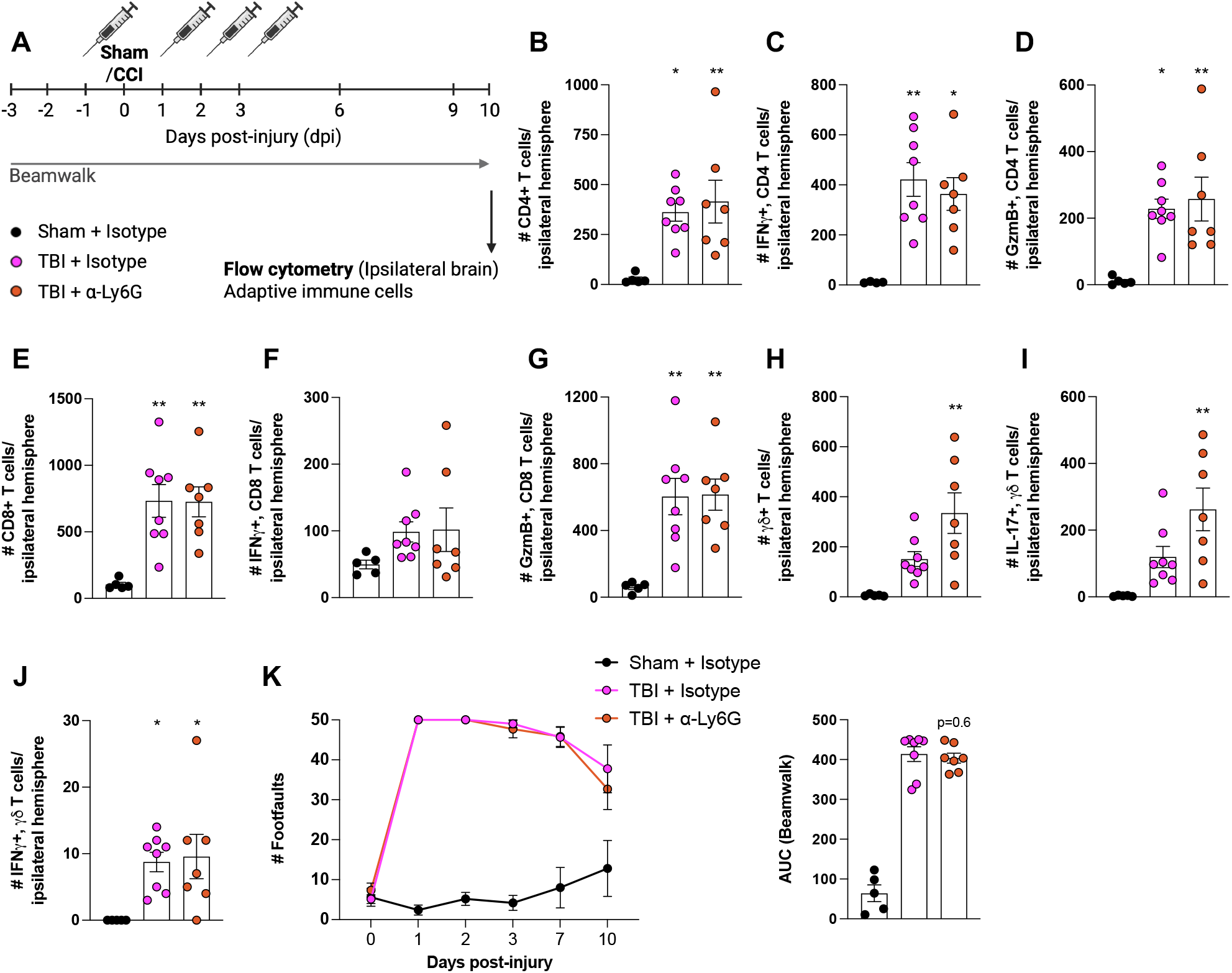
Neutrophil depletion does not alter T cell numbers or their effector phenotype and does not improve neurobehavioural outcomes. (A) Experimental design. (B-J) Bar graphs illustrate the absolute numbers of CD4^+^ T cells (B), IFN-γ-producing CD4^+^ T cells (C), GzmB-producing CD4^+^ T cells (D), CD8^+^ T cells (E), IFN-γ-producing CD8^+^ T cells (F), GzmB-producing CD8^+^ T cells (G), γδ^+^ T cells (H), IL-17-producing γδ^+^ T cells (I), and IFN-γ-producing γδ^+^ T cells (J). Neurobehavioral tests: Beam walk (K). Data = Mean ± SEM (n=10 per group); Sham vs. TBI *p<0.05, **p<0.01, ***p<0.001; TBI + Isotype vs. TBI + α-Ly6G ^+^p<0.05, ^++^p<0.01, ^+++^p<0.001 by one-way ANOVA with Tukey’s multiple comparisons test. Abbreviations: Hp, hippocampus.

To assess whether early neutrophil depletion affects functional recovery, we evaluated motor performance using established neurobehavioral tests. Fine motor coordination was assessed using a beamwalk test over the first 10 dpi. When compared to sham, there were robust TBI-induced deficits in motor function in isotype control treated TBI mice (**Fig. 4K**), but there were no difference in motor function performance in α-Ly6G treated TBI mice. The area-under-the-curve analysis indicated no statistically significant difference between the TBI + α-Ly6G and TBI + isotype groups (p=0.6). Overall, these data indicated that neutrophil depletion did not result in a sustained suppression of the sub-acute adaptive immune response following TBI. Instead, it was associated with a selective enhancement of the IL-17^+^-γδ^+^ T cell axis, without any observable improvements in neurobehavioural outcomes.

### Monocyte depletion suppresses the recruitment and effector activation of CD8^+^ and γδ^+^ T cell subsets in the injured brain

Monocytes constituted a significant secondary wave of IL-1β^+^ peripheral leukocyte recruitment to the injured brain following the initial neutrophil response. We therefore depleted inflammatory monocytes using MC-21, a function-blocking monoclonal antibody directed against CCR2 that depletes Ly6C^high^ monocytes in mice [19]. In proof-of-concept studies we administered MC-21 I.P. or an isotype control rat IgG2b antibody to mice 12h prior to TBI. Blood and brain were collected at 12h post-injury. This resulted in a complete depletion of monocytes in the circulation and injured brain when compared to the isotype control group (**Suppl. Fig. 6**). We then performed this monocyte depletion method in sham and CCI mice with repeated MC-21 treatment administered I.P. at day -1, 1 and 2 dpi. Rotarod assessment of gross motor function was performed prior to, and after TBI, and brain tissue was collected at 3 dpi for immune activation and flow cytometry analysis (**Fig. 5A**).

**Figure 5.**
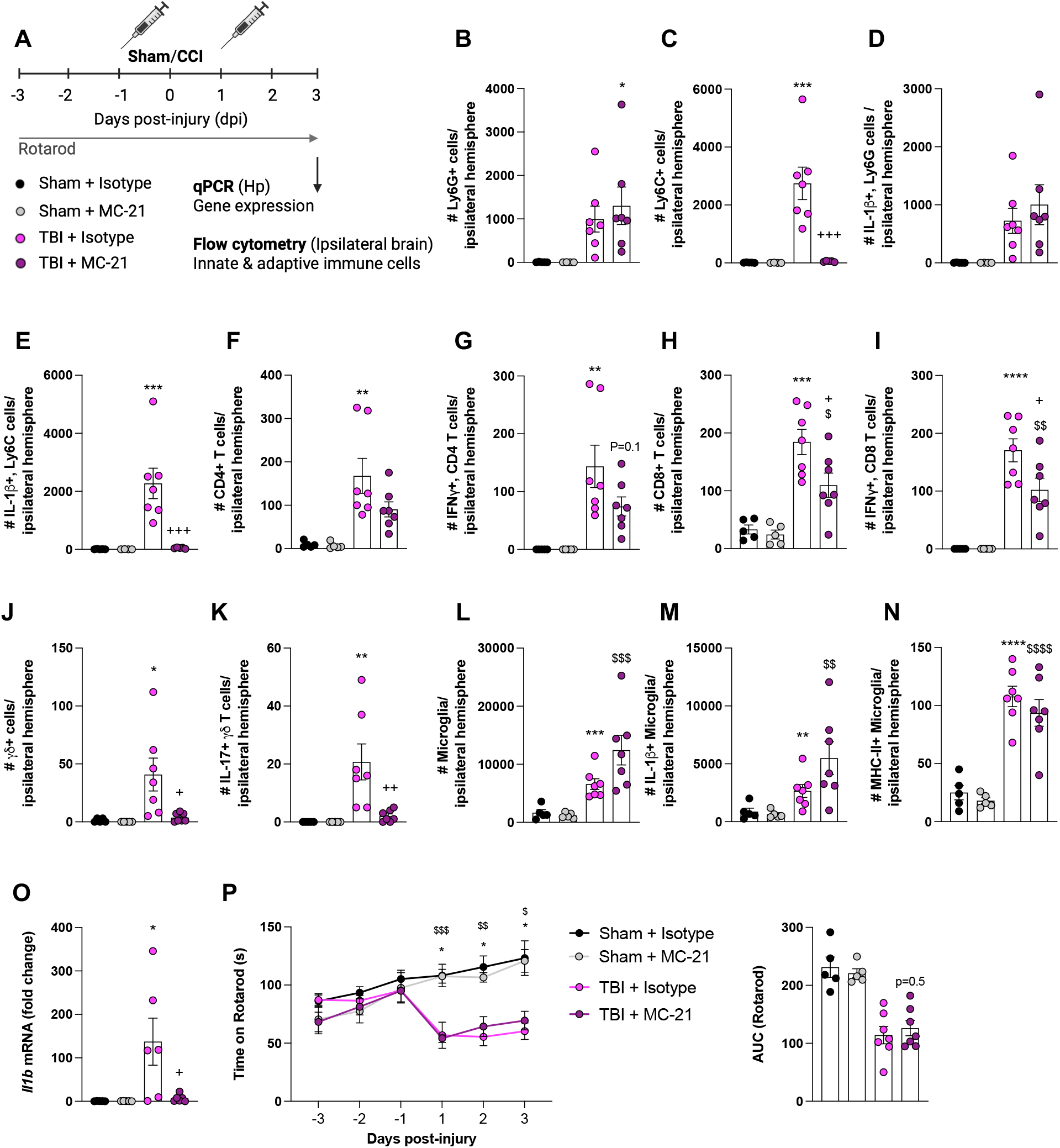
Monocyte depletion acutely alters T cell numbers and their effector phenotype post-TBI. (A) Experimental design. (B-N) Bar graphs illustrate the absolute numbers of Ly6G^+^ neutrophils (B), Ly6C^+^ monocytes (C), IL-1β-producing neutrophils (D), IL-1β-producing monocytes (E), CD4^+^ T cells (F), IFN-γ-producing CD4^+^ T cells (G), CD8^+^ T cells (H), IFN-γ-producing CD8^+^ T cells (I), γδ^+^ T cells (J), IL-17-producing γδ^+^ T cells (K), microglia (L), IL-1β-producing microglia (M), MHC-II^+^ microglia (N). Relative fold-change expression levels of *IL-1β* mRNA in hippocampus (O). Neurobehavioural tests: Rotarod (P). Data are presented as Mean ± SEM (n=5-7 per group); Sham + Isotype vs. TBI + Isotype *p<0.05, **p<0.01, ***p<0.001; Sham + MC-21 vs. TBI + MC-21 ^$^p<0.05, ^$$^p<0.01, ^$$$^p<0.001; TBI + Isotype vs. TBI + MC-21 ^+^p<0.05, ^++^p<0.01, ^+++^p<0.001 by two-way ANOVA with post-hoc Tukey’s test to correct for multiple comparisons. Abbreviations: Hp, hippocampus.

TBI increased the numbers of Ly6G^+^ neutrophils at 3 dpi (**Fig. 5B**), but MC-21 treatment did not mitigate the neutrophil accumulation. TBI induced a substantial increase in the numbers of Ly6C^+^ monocytes (**Fig. 5C**), which was completely abrogated by MC-21 treatment. The increase in IL-1β^+^monocytes following TBI was reduced to near-baseline levels with MC-21 treatment (**Fig. 5E**), whereas IL-1β^+^ neutrophils remained elevated with MC-21 treatment (**Fig. 5D**). Furthermore, monocyte depletion influenced early T cell responses after TBI. Total CD4^+^ T cells increased post-TBI (**Fig. 5F**), but MC-21 did not significantly reduce the numbers of CD4^+^ T cells, while there was a trend towards a reduction of IFN-γ^+^ CD4 T cells with MC-21 treatment (**Fig. 5G**). Notably, MC-21 treatment reduced the accumulation of CD8^+^ T cells in the injured brain (**Fig. 5H**), and also reduced IFN-γ^+^-CD8^+^ T cells when compared with the TBI + Isotype control group (**Fig. 5I**). Furthermore, MC-21 treatment reduced total γδ^+^ T cells after TBI (**Fig. 5J**), and IL-17^+^-γδ^+^ T cells when compared to the TBI + Isotype control group (**Fig. 5K**).

TBI increased the total numbers of microglia (**Fig. 5L**) and IL-1β^+^ microglia (**Fig. 5M**), while MC-21 treatment increased numbers of IL-1β^+^ microglia at 3 dpi, although this did not reach statistical significance (**Fig. 5L,M**). TBI also increased the numbers of MHC-II^+^ microglia (**Fig. 5N**), but MHC-II expression was not altered by MC-21 treatment. Consistent with the cellular findings, *Il1b* mRNA expression in the hippocampus was reduced by MC-21 treatment when compared to levels in the TBI + Isotype control group (**Fig. 5O**). Functionally, TBI-induced impairments in rotarod performance were not altered by monocyte depletion by MC-21 treatment (**Fig. 5P**). The area-under-the-curve analysis showing no significant difference between the TBI + MC-21 and TBI + isotype groups (p=0.5). These data indicate that CCR2^+^ inflammatory monocytes serve as crucial regulators of acute neuroinflammation after TBI, facilitating early IL-1β transcription and promoting both the recruitment and effector activation of CD8^+^ and γδ^+^ T cell subsets. Nevertheless, inflammatory monocyte depletion does not enhance early motor function recovery suggesting that monocyte-dependent early T cell modulation is insufficient to ameliorate acute neurological impairments in this model.

### Monocyte depletions results in partial motor function recovery but fails to alter T cell effector phenotypes at 10 dpi

We then investigated whether monocyte depletion influenced T cell responses and neurobehavioural outcomes up to 10 dpi (**Fig. 6A**). TBI resulted in a robust and sustained increase in total numbers of CD4^+^ T cells at 10 dpi (**Fig. 6B**), along with elevated levels of IFN-γ^+^- (**Fig. 6C**) and GzmB^+^- (**Fig. 6D**) CD4^+^ T cells. There was no reduction in CD4^+^ T cell numbers with MC-21 treatment (**Fig. 6B-D**). Similarly, total numbers of CD8^+^ T cells (**Fig. 6E**), IFN-γ^+^-(**Fig. 6F**) and GzmB^+^-(**Fig. 6G**) CD8^+^ T cells remained significantly elevated at 10 dpi. There was no effect of MC-21 treatment on the CD8 compartment at this later time point (**Fig. 6E-G**). Finally, total γδ^+^ T cells (**Fig. 6H**) were increased at 10 dpi, as were IL-17^+^- (**Fig. 6I**) and IFN-γ^+^- (**Fig. 6J**) γδ^+^ T cells, but MC-21 treatment failed to reduce γδ^+^ T cells numbers (**Fig. 6H-J**).

**Figure 6.**
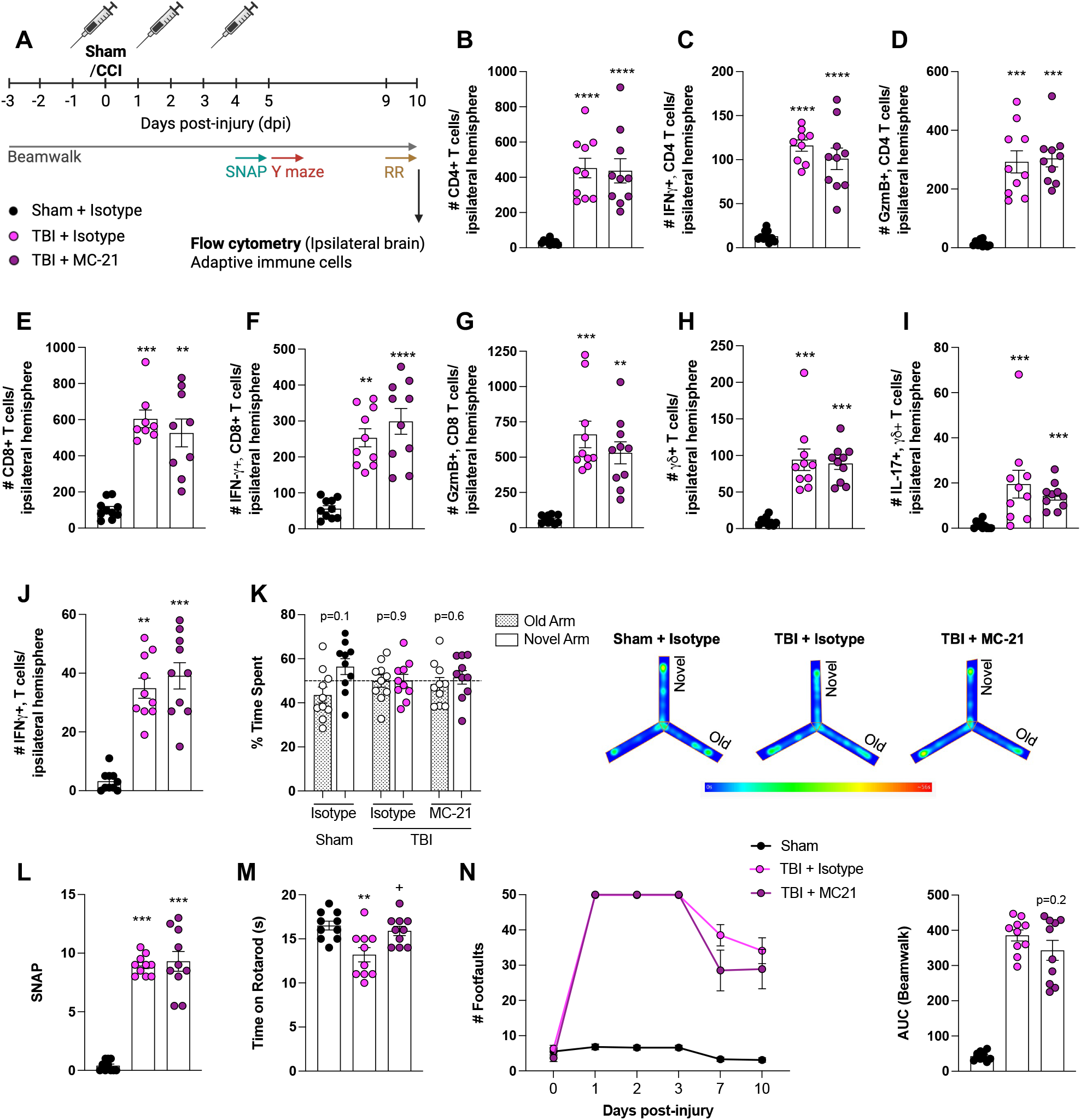
Partial motor function recovery is observed at 10 dpi following monocyte depletion despite persistent T cell infiltration. (A) Experimental design. (B-J) Bar graphs illustrate the absolute numbers of CD4^+^ T cells (B), IFN-γ-producing CD4^+^ T cells (C), GzmB-producing CD4^+^ T cells (D), CD8^+^ T cells (E), IFN-γ-producing CD8^+^ T cells (F), GzmB-producing CD8+ T cells (G), γδ^+^ T cells (H), IL-17-producing γδ^+^ T cells (I), and IFN-γ-producing γδ^+^ T cells (J). (K-N) Neurobehavioral tests: Y-maze (K), SNAP score (L), Forced rotarod (M), Beam walk (N). Data = Mean ± SEM (n=10 per group); Sham vs. TBI *p<0.05, **p<0.01, ***p<0.001; TBI + Isotype vs. TBI + MC-21 ^+^p<0.05, ^++^p<0.01 by one-way ANOVA with Tukey’s multiple comparisons test. Abbreviations: Hp, hippocampus.

To assess whether monocyte depletion affected functional recovery, we evaluated cognitive and motor performance using established neurobehavioral tests. In the 2-trial Y-maze task, sham mice spent more time in the novel arm compared to the familiar arm, suggesting intact short-term spatial working memory; however, this discrimination did not achieve statistical significance (paired t-test, p = 0.1; **Fig.6K**). TBI induced a clear spatial memory deficit, that was not rescued by MC-21 treatment (paired t-test, TBI + isotype, p = 0.9; TBI + MC-21, p = 0.6). Similarly, MC-21 treatment failed to attenuate the TBI-induced sensorimotor impairment assessed by the SNAP test (p<0.001 vs. sham; **Fig. 6L**). In contrast, in a once off accelerating rotarod test performed at 10 dpi MC-21 treated TBI mice showed improved motor performance and spent significantly more time on the rotarod compared to TBI mice treated with isotype control (p<0.05 vs. TBI + Isotype; **Fig. 6M**). Fine motor coordination was assessed using a beamwalk test over the first 10 dpi. TBI induced deficits in motor coordination in both MC-21- and isotype-treated TBI mice up to 3 dpi compared to sham (**Fig.6N**). The partial spontaneous recovery in motor function at 7 and 10 dpi was further improved by MC-21 treatment, although it did not reach statistical significance, as indicated by the area-under-the-curve analysis (p=0.2). Overall, the data demonstrate that MC-21 mediated monocyte depletion did not alter the T cell response in the sub-acute phase post-injury. However, monocyte depletion resulted in subtle improvements in neurological outcomes, particularly related to TBI-induced persistent deficits in fine motor function.

### NLRP3 inflammasome inhibition reduces microglial IL-1β without altering T cell effector phenotype

Based on modest effects of neutrophil and monocyte depletion on T cell responses after TBI, we hypothesized that inflammasome-dependent IL-1β signalling via NLRP3 may be an upstream mechanism influencing the post-injury adaptive immune environment. To investigate this, we used MCC950, a selective NLRP3 inflammasome inhibitor [20]. In pilot therapeutic studies we administered MCC950 (10mg/kg, I.P.) or Vehicle (saline) starting at 2 hpi with repeated dosing at 12, 24, and 48 hpi, and brain tissue was collected at 3 dpi. MCC950 treatment resulted in a strong suppression of IL-1β^+^ microglia after TBI (**Suppl. Fig. 7**), along with non-significant reductions in IL-1β^+^ infiltrating monocytes and neutrophils. We repeated the MCC950 treatment protocol in sham and CCI mice, and rotarod assessment of gross motor function was performed prior to, and after TBI, and brain tissue was collected at 3 dpi for immune activation and flow cytometry analysis (**Fig. 7A**).

**Figure 7.**
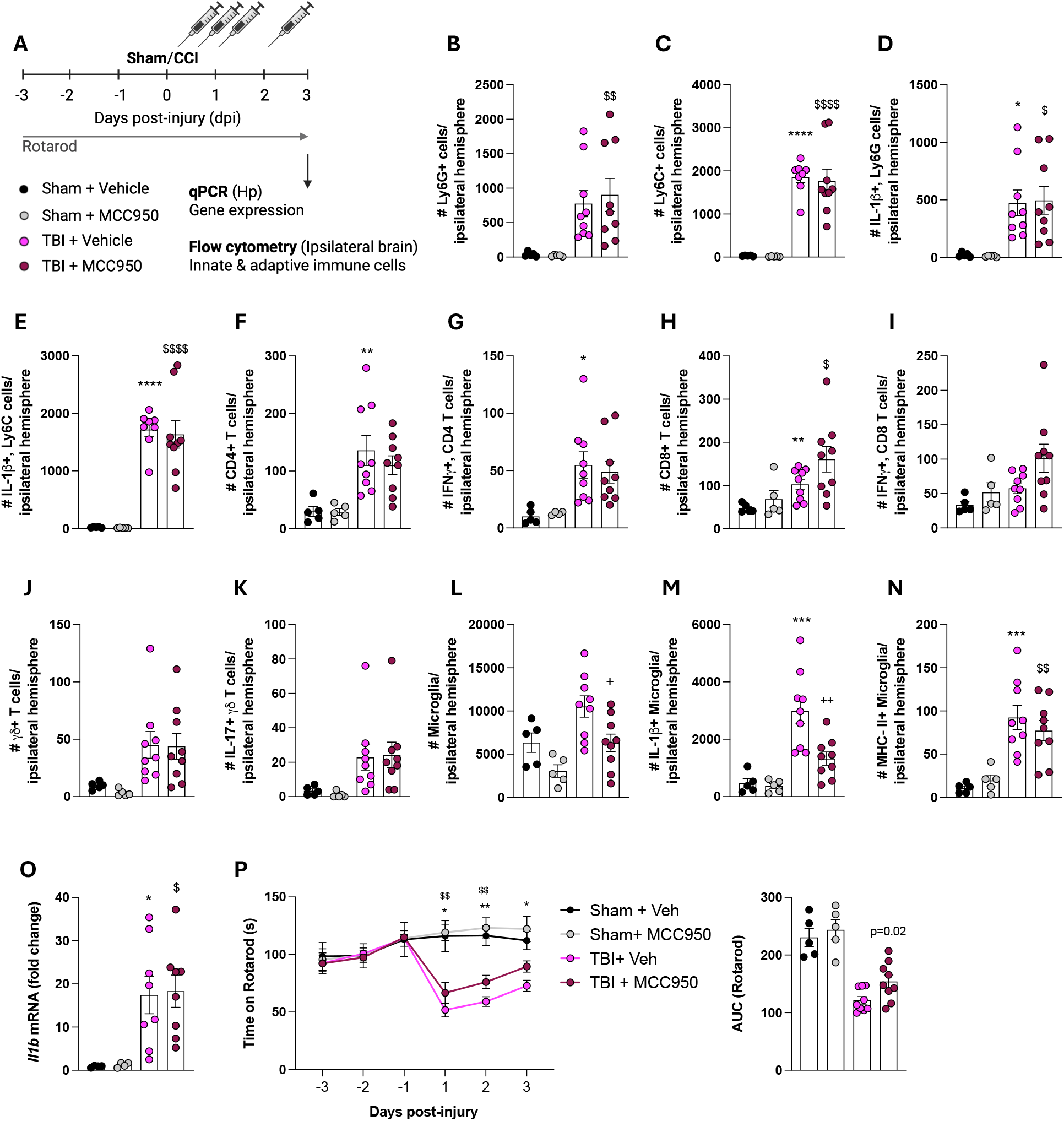
MCC950 treatment selectively reduces microglial IL-1β at 3dpi without altering T cell effector phenotype. (A) Experimental design. (B-N) Bar graphs illustrate the absolute numbers of Ly6G^+^ neutrophils (B), Ly6C^+^ monocytes (C), IL-1β-producing neutrophils (D), IL-1β-producing monocytes (E), CD4^+^ T cells (F), IFN-γ-producing CD4^+^ T cells (G), CD8^+^ T cells (H), IFN-γ-producing CD8^+^ T cells (I), γδ^+^ T cells (J), IL-17-producing γδ^+^ T cells (K), microglia (L), IL-1β-producing microglia (M), and MHC-II^+^ microglia (N). Relative fold-change expression levels of *Il1b* mRNA in hippocampus (O). Neurobehavioural tests: Rotarod (P). Data are presented as Mean ± SEM (n=5-9 per group); Sham + Vehicle vs. TBI + Vehicle *p<0.05, **p<0.01, ***p<0.001; Sham + MCC950 vs. TBI + MCC950 ^$^p<0.05, ^$$^p<0.01, ^$$$^p<0.001; TBI + Vehicle vs. TBI + MCC950 ^+^p<0.05, ^++^p<0.01, ^+++^p<0.001 by two-way ANOVA with post-hoc Tukey’s test to correct for multiple comparisons. Abbreviations: Hp, hippocampus.

TBI increased the total numbers of infiltrating Ly6G^+^ neutrophils (**Fig. 7B**) and Ly6C^+^ monocytes (**Fig. 7C**) at 3 dpi, as well as IL-1β^+^ neutrophils (**Fig. 7D**) and IL-1β^+^ monocytes (**Fig. 7E**) when compared to sham. However, MCC950 treatment did not reduce the accumulation of these infiltrating myeloid populations (**Fig. 7B-E**). Within the T cell compartment, TBI increased the total numbers of CD4^+^ (**Fig. 7F**) and CD8^+^ (**Fig. 7H**) T cells, as well as IFN-γ^+^-CD4^+^ (**Fig. 7G**) and CD8^+^ (**Fig. 7I**) T cells when compared to sham. MCC950 treatment failed to reduce T cell numbers at 3 dpi. Notably, there was a non-significant trend towards increased IFN-γ^+^-CD8^+^ T cell expression in the injured brain following NLRP3 inflammasome inhibition (**Fig. 7I**). While total γδ^+^ T cells (**Fig. 7J**) and IL-17^+^-γδ^+^ T cells (**Fig. 7K**) were increased after TBI, they were not significantly altered by MCC950 treatment (**Fig. 7J,K**).

In contrast to the limited effects on infiltrating leukocyte numbers, MCC950 treatment robustly altered the resident innate immune compartment (i.e. microglia). NLRP3 inflammasome inhibition by MCC950 reduced the TBI-induced increase in total microglia (**Fig. 7L**), IL-1β^+^ microglia (**Fig. 7M**), and MHC-II^+^ microglia (**Fig. 7N**); although the latter did not reach statistical significance. At the tissue level, despite effects of MCC950 treatment on microglia there was no difference in hippocampal *Il1b* mRNA expression between vehicle-treated and MCC950-treated TBI mice (**Fig. 7O**). Functionally, TBI-induced impairments in rotarod performance were not rescued by MCC950 treatment (**Fig. 7P**). However, area-under-the-curve analysis revealed a significant improvements due to MCC950 treatment (p=0.02), indicating an overall benefit in motor performance over the acute post-injury testing period. These data demonstrate that inhibiting NLRP3 inflammasome with MCC950 had a negligible effect on early leukocyte infiltration and IL-1β expressing peripheral myeloid subsets following TBI. Nonetheless, MCC950 treatment specifically and robustly reduced microglial activation, including IL-1β^+^ microglia, which was associated with a modest improvements in motor function in the early post-injury days.

### NLRP3 inflammasome inhibition improves motor and cognitive function after TBI without altering T cell effector phenotypes

We finally investigated whether NLRP3 inflammasome inhibition could influence T cells and neurobehavioural outcomes up to 10 dpi (**Fig. 8A**). TBI resulted in a robust and sustained increase in the total numbers of CD4^+^ T cells (**Fig. 8B**) at 10 dpi, along with elevated levels of IFN-γ^+^- (**Fig. 8C**) and GzmB^+^- (**Fig. 8D**) CD4^+^ T cells. There was no reduction in CD4^+^ T cell numbers with MCC950 treatment (**Fig. 8B-D**). Similarly, total numbers of CD8^+^ T cells (**Fig. 8E**), IFN-γ^+^- (**Fig. 8F**) and GzmB^+^- (**Fig. 8G**) CD8^+^ T cells remained elevated at 10 dpi. There was a minor reduction in the GzmB^+^-CD8^+^ T cells with MCC950 treatment (**Fig. 8G**), but this failed to reach statistical significance. There was no effect of treatment on IFN-γ^+^ T cells (**Fig. 8F**). Total γδ+ T cells (**Fig. 8H**) were increased at 10 dpi, as were IL-17^+^- (**Fig. 8I**) and IFN-γ^+^- (**Fig. 8J**) γδ^+^ T cells. However, MCC950 treatment failed to reduce γδ^+^ T cells numbers (**Fig. 8H-J**).

**Figure 8.**
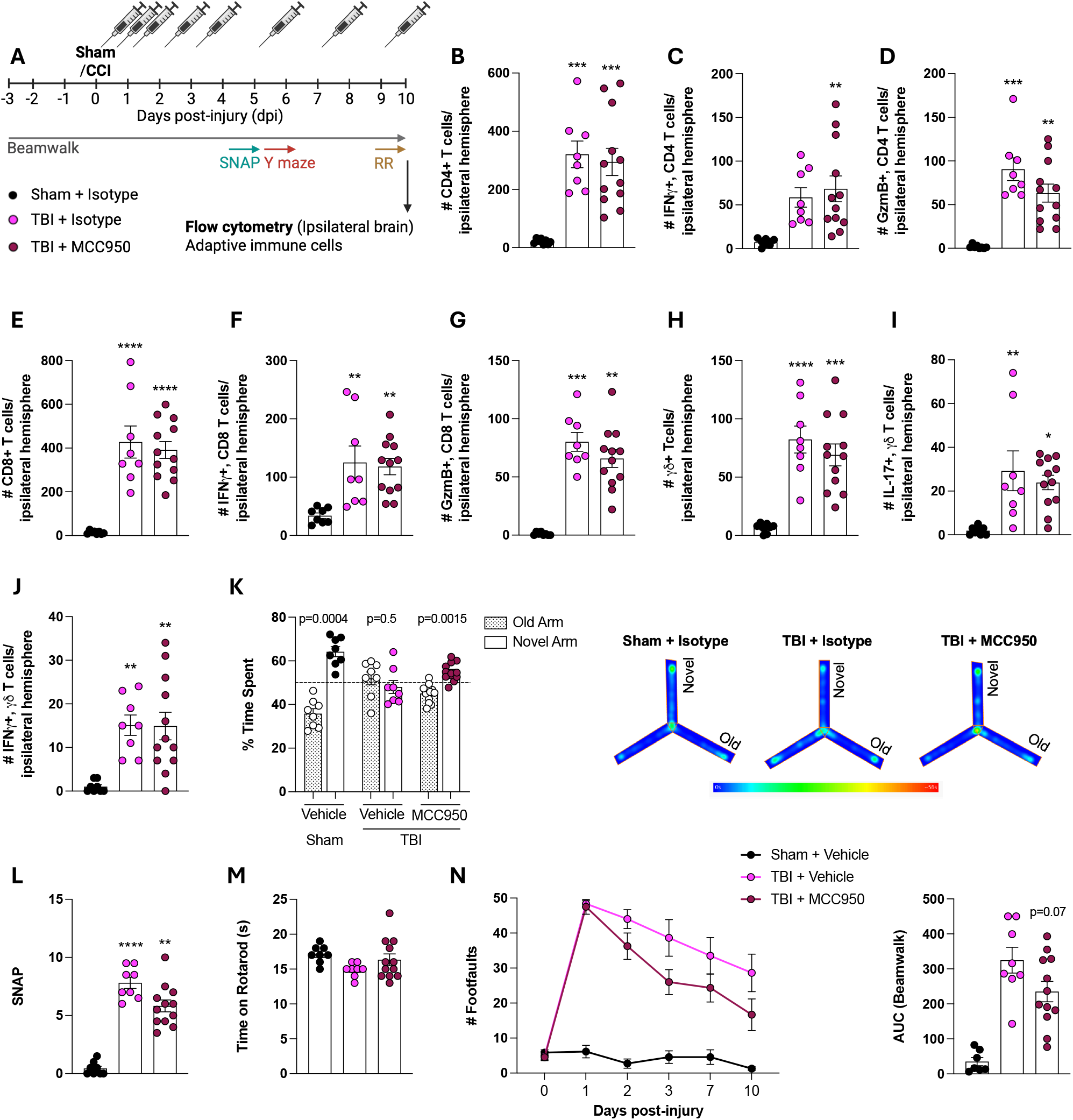
MCC950 does not alter T cell effector phenotype but had beneficial effects on motor and cognitive function at 10 dpi. (A) Experimental design. (B-J) Bar graphs illustrate the absolute numbers of CD4^+^ T cells (B), IFN-γ-producing CD4^+^ T cells (C), GzmB-producing CD4^+^ T cells (D), CD8^+^ T cells (E), IFN-γ-producing CD8^+^ T cells (F), GzmB-producing CD8^+^ T cells (G), γδ^+^ T cells (H), IL-17-producing γδ^+^ T cells (I), IFN-γ-producing γδ+ T cells (J). (K-N) Neurobehavioral tests: Y-maze (K), SNAP score (L), Forced rotarod (M), Beam walk (N). Data= Mean ± SEM (n=8-12 per group); Sham + Vehicle vs. TBI + Vehicle *p<0.05, **p<0.01,***p<0.001 by one-way ANOVA with Tukey’s multiple comparisons test. Abbreviations: Hp, hippocampus.

To determine whether NLRP3 inflammasome inhibition could improve functional recovery after TBI, we evaluated cognitive and motor performance using established neurobehavioral tests. In the 2-trial Y maze task, sham + vehicle mice spent more time in the novel arm compared to the familiar arm, showing a clear novel-arm preference (paired t-test, p=0.0004). In contrast, TBI + vehicle mice had cognitive deficits and did not show a preference (paired t-test, p=0.5), while MCC950-treated TBI mice spent more time in the novel arm demonstrating novel-arm preference (paired t-test, p=0.0015), indicating improved cognitive performance (**Fig. 8K**). Similarly, SNAP scoring revealed TBI-induced deficits in gross motor function (p<0.001 vs. sham; **Fig. 8L**), while there was a trend to improved motor function in the MCC950-treated group. In a once off accelerating rotarod test performed at 10 dpi vehicle treated TBI mice spent less time on the rotarod when compared to sham (p=0.1 vs. sham; **Fig. 8M**), while MCC950 treated TBI mice had improved motor performance and spent more time on the rotarod (p=0.3 vs. Vehicle + TBI). In the beamwalk test, there were robust TBI-induced deficits in fine motor coordination in vehicle treated TBI mice when compared to sham (**Fig. 8N**). There were improvements in footfault scores in MCC950 treated TBI mice at 3 and 10 dpi (**Fig. 8N**), as indicated by the area-under-the-curve analysis (p=0.07). Overall, MCC950-mediated NLRP3 inflammasome inhibition did not alter the T cell response in the sub-acute phase up to 10 dpi. However, it resulted in subtle improvements in neurological outcomes, particularly related to TBI induced persistent deficits in fine motor coordination and spatial learning.

## Discussion

TBI initiates a sequential immune response characterized by the early recruitment of neutrophils and CCR2-dependent monocytes, which culminates in sustained microglial activation and prolonged T cell infiltration [21]. Building upon this framework, our central hypothesis is that the primary early cellular source of IL-1β promotes subsequent effector T cell responses, particularly IL-17- and IFN-γ-secreting T cells, which contribute to behavioural dysfunction. IL-1 signalling is an established driver of Th_17_ programming and IL-17 production, including through direct T cell IL-1 receptor engagement [22]. Additionally, IL-1 family cytokines serve as instructors and amplifiers of type-1 effector function, supporting IFN-γ production from T cells under inflammatory conditions [23]. Notably, emerging TBI literature positions IL-17- and IFN-γ-producing T cell responses as precursors to maladaptive neuroinflammatory amplification, including microglial activation [24, 25].

Our study examines whether early innate perturbation (α-Ly6G and MC-21) or inhibition of the upstream pathway (NLRP3 inflammasome via MCC950) can reprogram subsequent adaptive immune responses and improve behavioural outcomes. We show that each intervention resulted in biologically significant alterations in immune responses; however, the neurological benefits were modest and specific to certain domains. This aligns with the redundancy observed in post-TBI inflammatory processes. Antibody-depletion based targeting of neutrophils altered the early inflammatory composition but did not result in a sustained reprogramming of subacute neuroimmune cellular responses or behaviour. Moreover, it enhanced IL-17 production by γδ^+^T cells and IFN-γ by CD4^+^ and CD8^+^ T cells that were sustained through 10 dpi. This is consistent with reports showing that neutrophils restrain IL-17-producing γδ^+^ T cells through a contact dependent mechanism, such that neutrophil depletion can release this brake and amplify IL-17 responses [26]. Neutrophils are recognized as early amplifiers of secondary injury after TBI, and their depletion or impaired recruitment can mitigate edema and tissue pathology [27]. However, functional recovery often remains relatively unaffected, as has been demonstrated in CCI and closed head injury models, including those with CXCR2 disruption [28]. CCR2^high+^ monocyte depletion by MC-21 influenced early programming but did not durably suppress subacute inflammation. While MC-21 treatment robustly reduced TBI-induced expression of IL-1β by monocytes, IFN-γ by CD4^+^ and CD8^+^ T cells, and IL-17 by γδ^+^ T cells at 3 dpi, it also increased IL-1β production in microglia. CCR2-dependent monocytes and monocyte-derived macrophages contribute to post-TBI inflammatory polarization and cognitive impairments [29, 30], while CCR2 antagonism (using CCX872) reduces inflammatory macrophage accumulation and improves cognition, supporting a causal CCR2 inflammatory axis [30]. We found that early monocyte depletion did not consistently attenuate inflammation by 10 dpi, likely due to compensatory recruitment and adaptation of resident glia because CCR2-linked monocyte availability shapes microglial subset trajectories such that reduced monocyte influx can shift inflammatory execution towards brain resident microglia rather than lowering the total monocyte-mediated inflammatory burden [31].

In comparison to depletion strategies that focus on a single cellular compartment, the behavioural sensitivity to MCC950 observed in our preclinical TBI modelling aligns with the decoupling of leukocyte recruitment from inflammasome-associated IL-1β output. We determined that MCC950 specifically and robustly attenuated IL-1β production in microglia, while leukocyte recruitment remained largely unaffected. This pattern supports the hypothesis that inhibiting an upstream inflammasome node can significantly reduce the parenchymal IL-1β signal by limiting the microglial contribution, even when peripheral myeloid cells continue to be recruited. Prior TBI studies showed that NLRP3 inhibition with MCC950 enhances neurological outcomes, decreases edema and lesion volume, while also suppressing pro-inflammatory gene/protein expression patterns in injured tissues [6, 8]. When combined, our findings suggest that recruitment alone is insufficient to sustain IL-1β signalling if microglial inflammasome activity, a major local amplifier, is suppressed. Thus, compensatory cellular immune remodelling is identified as a mechanistic constraint rather than a secondary observation. Studies have shown that suppressing one myeloid compartment can lead to the redistribution of effector functions to another, rather than diminishing the overall inflammatory capacity [32, 33]. In CNS disease/injury contexts, reciprocal interactions between microglia and macrophages have been demonstrated to adjust microglial inflammatory programs and phagocytic behaviour when the availability of infiltrating monocyte-derived macrophages is altered [34]. A similar principle is observed in white matter demyelination and repair biology, where the depletion of microglia does not impede myelin debris clearance because monocyte-derived macrophages can compensate, thereby maintaining clearance while altering the executing compartment [35]. From this perspective, the increase in monocytes following neutrophil depletion and the increase in microglia following monocyte depletion are best understood as network-level reallocation of the IL-1β competent myeloid capacity. This is relevant to our hypothesis because such source switching can sustain the early IL-1β environment that predisposes subsequent IL-17 and IFN-γ T cell responses, thereby supporting why shifts in immune composition may not correlate with neurobehavioural recovery.

There are limitations in our studies that need to be discussed. Specifically, our multispectral flow cytometry immunophenotyping protocols were optimized for T cell cytokine readouts and may therefore be sub-optimal for the diversity of microglial phenotypes and functions; albeit microglial IL-1β, MCH-II and CD68 were consistently measured by our protocols. The network effects are further influenced by the practical limitations of antibody-based depletion, which is frequently incomplete and challenging to sustain over time. This complicates the differentiation between true pathway suppression and rebound or replacement phenomena. The validation of α-Ly6G through flow cytometry is particularly challenging because antibody binding can obscure epitopes and alter perceived marker intensity, with efficacy varying according to context [36, 37]. Similarly, MC-21 is most effective within short dosing intervals due to the potential development of neutralizing anti-MC-21 antibodies, which can diminish monocyte depletion efficacy over time [13]. Consequently, null effects observed at later time points may be indicative of compensatory mechanisms and declining antibody performance, rather than a true disengagement of the targeted pathway. Additionally, the use of male-only cohorts and a limited therapeutic window may collectively reduce how well these findings generalize and how precisely they explain mechanism, particularly in females, who often show lower inflammatory cytokine responses after TBI [15].

Finally, T cell responses following TBI are increasingly recognized as a persistent, context-dependent contributor to chronic neuroinflammation, rather than a transient bystander. Chronic studies using CCI models demonstrate the sustained presence of CD4^+^ and CD8^+^ T cells [24, 38] for months post-injury accompanied by persistent gliosis [39]. Moreover, pharmacological depletion of CD8^+^ T cells, but not depletion of CD4^+^ T cells, improves long-term neurological outcomes in TBI mice and produces a neuroprotective Th_2_/Th_17_ immunological shift, indicating a persistent detrimental role for cytotoxic T cells in the chronically injured brain [24]. These studies indicate that restricting immune cell trafficking can mitigate chronic accumulation of macrophages and lymphocytes with sex-dependent functional protection [40]. However, the evidence is somewhat conflicting in the acute phase post-injury. For example, genetic ablation of lymphocytes does not confer neuroprotection immediately after TBI, as shown in Rag1-/- mice that have no early neuroprotection following closed head injury [41]. Additionally, modulation of the sphingosine-1-phosphate receptor with FTY720 reduces circulating lymphocytes and brain invasion without improving lesion size or behavioural outcomes [42]. However, another study showed that FTY720 partially improves neurobehavioural recovery, reduces brain edema as well as apoptotic cell death after TBI [43]. Our study extends this research by investigating a more mechanistic, hypothesis-driven question, whether early innate perturbations that modify the IL-1β axis can reprogram the subsequent T cell phenotype (rather than merely reducing immune presence), thereby influencing chronic microglial states and behaviour. Evidence supporting the concept that the quality of T cells can direct microglial activity is provided by intranasal anti-CD3, which induces IL-10 dependent FoxP3^+^ regulatory T cells that engage microglia, enhance phagocytosis, and improve functional recovery [38]. Therefore, it suggests that the timing and cellular source of early IL-1β could alter later T cell responses, biasing it toward pathogenic IL-17/IFN-γ-producing T cell subsets, thereby shaping chronic microglial trajectories and functional outcomes. A crucial next step involves the temporal mapping and source-specific manipulation of IL-1β-producing cells that will require advanced genetic access and lineage-tracing techniques. Such approaches may identify the initial IL-1β sources that are essential for initiating T cell polarization associated with secondary injury progression and recovery following TBI [44].

## Acknowledgments

We thank Professor Kingston Mills’ research team for critical feedback and discussion. In particular, we thank Dr. Barry Moran for guidance and advice with flow cytometry in the School of Biochemistry & Immunology, Trinity College Dublin.

## References

1. Maas AIR, Menon DK, Manley GT, Abrams M, Åkerlund C, Andelic N, Aries M, Bashford T, Bell MJ, Bodien YG, et al: Traumatic brain injury: progress and challenges in prevention, clinical care, and research. Lancet Neurol 2022, 21:1004–1060.

2. Simon DW, McGeachy MJ, Bayır H, Clark RS, Loane DJ, Kochanek PM: The far-reaching scope of neuroinflammation after traumatic brain injury. Nat Rev Neurol 2017, 13:171–191.

3. Faden AI, Loane DJ: Chronic neurodegeneration after traumatic brain injury: Alzheimer disease, chronic traumatic encephalopathy, or persistent neuroinflammation? Neurotherapeutics 2015, 12:143–150.

4. Lindblad C, Rostami E, Helmy A: Interleukin-1 Receptor Antagonist as Therapy for Traumatic Brain Injury. Neurotherapeutics 2023, 20:1508–1528.

5. Ravula AR, Murray KE, Rao KVR, Pfister BJ, Citron BA, Chandra N: MCC950 Attenuates Microglial NLRP3-Mediated Chronic Neuroinflammation and Memory Impairment in a Rat Model of Repeated Low-Level Blast Exposure. J Neurotrauma 2024, 41:1450–1468.

6. Xu X, Yin D, Ren H, Gao W, Li F, Sun D, Wu Y, Zhou S, Lyu L, Yang M, et al: Selective NLRP3 inflammasome inhibitor reduces neuroinflammation and improves long-term neurological outcomes in a murine model of traumatic brain injury. Neurobiol Dis 2018, 117:15–27.

7. Placeres-Uray F, Gorthy AS, Torres MD, Atkins CM: Inhibition of microglia priming by NLRP3 reduces the impact of early life stress and mild TBI. J Neuroinflammation 2025, 22:185.

8. Ismael S, Nasoohi S, Ishrat T: MCC950, the Selective Inhibitor of Nucleotide Oligomerization Domain-Like Receptor Protein-3 Inflammasome, Protects Mice against Traumatic Brain Injury. J Neurotrauma 2018, 35:1294–1303.

9. O’Brien WT, Pham L, Symons GF, Monif M, Shultz SR, McDonald SJ: The NLRP3 inflammasome in traumatic brain injury: potential as a biomarker and therapeutic target. J Neuroinflammation 2020, 17:104.

10. Kilgore MD, Xiu Y, Jiang Y, Wang Y, Shi M, Zhou D, Sein T, Vodovoz SJ, Wang D, Dumont AS, et al: T Cell Involvement in Neuroinflammation After Traumatic Brain Injury: Implications for Therapeutic Intervention. CNS Neurosci Ther 2025, 31:e70580.

11. Laabei J, Vegliante G, Strogulski NR, Douglas C, Threja S, Pearson A, Nkiliza A, Hanscom M, Dias Filogonio Emediato I, Crawford F, et al: The NOX2-ROS-NLRP3 inflammasome axis in traumatic brain injury. J Neuroinflammation 2025, 22:242.

12. Boivin G, Faget J, Ancey PB, Gkasti A, Mussard J, Engblom C, Pfirschke C, Contat C, Pascual J, Vazquez J, et al: Durable and controlled depletion of neutrophils in mice. Nat Commun 2020, 11:2762.

13. Lösslein AK, Lohrmann F, Scheuermann L, Gharun K, Neuber J, Kolter J, Forde AJ, Kleimeyer C, Poh YY, Mack M, et al: Monocyte progenitors give rise to multinucleated giant cells. Nat Commun 2021, 12:2027.

14. Hanscom M, Loane DJ, Aubretch T, Leser J, Molesworth K, Hedgekar N, Ritzel RM, Abulwerdi G, Shea-Donohue T, Faden AI: Acute colitis during chronic experimental traumatic brain injury in mice induces dysautonomia and persistent extraintestinal, systemic, and CNS inflammation with exacerbated neurological deficits. J Neuroinflammation 2021, 18:24.

15. Doran SJ, Ritzel RM, Glaser EP, Henry RJ, Faden AI, Loane DJ: Sex Differences in Acute Neuroinflammation after Experimental Traumatic Brain Injury Are Mediated by Infiltrating Myeloid Cells. J Neurotrauma 2019, 36:1040–1053.

16. Vegliante G, Pischiutta F, Restelli E, Moro F, Chiaravalloti MA, Raimondi I, Bertani I, Lisi I, Sammali E, Pascente R, et al: Human Brain Contusions Contain Pathogenic Transmissible Species that Induce Progressive Cognitive Decline and Tau Pathology in Mice. Ann Neurol 2025 (online ahead of print).

17. Moro F, Pischiutta F, Portet A, Needham EJ, Norton EJ, Ferdinand JR, Vegliante G, Sammali E, Pascente R, Caruso E, et al: Ageing is associated with maladaptive immune response and worse outcome after traumatic brain injury. Brain Commun 2022, 4:fcac036.

18. Henry RJ, Ritzel RM, Barrett JP, Doran SJ, Jiao Y, Leach JB, Szeto GL, Wu J, Stoica BA, Faden AI, Loane DJ: Microglial Depletion with CSF1R Inhibitor During Chronic Phase of Experimental Traumatic Brain Injury Reduces Neurodegeneration and Neurological Deficits. J Neurosci 2020, 40:2960–2974.

19. Mack M, Cihak J, Simonis C, Luckow B, Proudfoot AE, Plachý J, Brühl H, Frink M, Anders HJ, Vielhauer V, et al: Expression and characterization of the chemokine receptors CCR2 and CCR5 in mice. J Immunol 2001, 166:4697–4704.

20. Coll RC, Robertson AA, Chae JJ, Higgins SC, Muñoz-Planillo R, Inserra MC, Vetter I, Dungan LS, Monks BG, Stutz A, et al: A small-molecule inhibitor of the NLRP3 inflammasome for the treatment of inflammatory diseases. Nat Med 2015, 21:248–255.

21. Jin X, Ishii H, Bai Z, Itokazu T, Yamashita T: Temporal changes in cell marker expression and cellular infiltration in a controlled cortical impact model in adult male C57BL/6 mice. PLoS One 2012, 7:e41892.

22. Chung Y, Chang SH, Martinez GJ, Yang XO, Nurieva R, Kang HS, Ma L, Watowich SS, Jetten AM, Tian Q, Dong C: Critical regulation of early Th17 cell differentiation by interleukin-1 signaling. Immunity 2009, 30:576–587.

23. Garlanda C, Dinarello CA, Mantovani A: The interleukin-1 family: back to the future. Immunity 2013, 39:1003–1018.

24. Daglas M, Draxler DF, Ho H, McCutcheon F, Galle A, Au AE, Larsson P, Gregory J, Alderuccio F, Sashindranath M, Medcalf RL: Activated CD8(+) T Cells Cause Long-Term Neurological Impairment after Traumatic Brain Injury in Mice. Cell Rep 2019, 29:1178–1191.e1176.

25. Abou-El-Hassan H, Rezende RM, Izzy S, Gabriely G, Yahya T, Tatematsu BK, Habashy KJ, Lopes JR, de Oliveira GLV, Maghzi AH, et al: Vγ1 and Vγ4 gamma-delta T cells play opposing roles in the immunopathology of traumatic brain injury in males. Nat Commun 2023, 14:4286.

26. Yu X, Yue X, Tchudjin Magatsin JD, Marwitz S, Behrends J, Goldmann T, Opferman JT, Kasper B, Petersen F: Neutrophils negatively control IL-17A-producing γδ T cell frequencies in a contact-dependent manner under physiological conditions. Front Immunol 2025, 16:1542191.

27. Kenne E, Erlandsson A, Lindbom L, Hillered L, Clausen F: Neutrophil depletion reduces edema formation and tissue loss following traumatic brain injury in mice. J Neuroinflammation 2012, 9:17.

28. Semple BD, Bye N, Ziebell JM, Morganti-Kossmann MC: Deficiency of the chemokine receptor CXCR2 attenuates neutrophil infiltration and cortical damage following closed head injury. Neurobiol Dis 2010, 40:394–403.

29. Paladini MS, Yang BA, Torkenczy KA, Frias ES, Feng X, Krukowski K, Sit R, Morri M, Lam W, Pedoia V, et al: Fate mapping of peripherally-derived macrophages after traumatic brain injury in mice reveals a long-lasting population with a distinct transcriptomic signature. Nat Commun 2025, 16:8898.

30. Morganti JM, Jopson TD, Liu S, Riparip LK, Guandique CK, Gupta N, Ferguson AR, Rosi S: CCR2 antagonism alters brain macrophage polarization and ameliorates cognitive dysfunction induced by traumatic brain injury. J Neurosci 2015, 35:748–760.

31. Somebang K, Rudolph J, Imhof I, Li L, Niemi EC, Shigenaga J, Tran H, Gill TM, Lo I, Zabel BA, et al: CCR2 deficiency alters activation of microglia subsets in traumatic brain injury. Cell Rep 2021, 36:109727.

32. Soehnlein O, Lindbom L: Phagocyte partnership during the onset and resolution of inflammation. Nat Rev Immunol 2010, 10:427–439.

33. Makinde HM, Cuda CM, Just TB, Perlman HR, Schwulst SJ: Nonclassical Monocytes Mediate Secondary Injury, Neurocognitive Outcome, and Neutrophil Infiltration after Traumatic Brain Injury. J Immunol 2017, 199:3583–3591.

34. Greenhalgh AD, Zarruk JG, Healy LM, Baskar Jesudasan SJ, Jhelum P, Salmon CK, Formanek A, Russo MV, Antel JP, McGavern DB, et al: Peripherally derived macrophages modulate microglial function to reduce inflammation after CNS injury. PLoS Biol 2018, 16:e2005264.

35. Baaklini CS, Ho MFS, Lange T, Hammond BP, Panda SP, Zirngibl M, Zia S, Himmelsbach K, Rana H, Phillips B, et al: Microglia promote remyelination independent of their role in clearing myelin debris. Cell Rep 2023, 42:113574.

36. Pollenus E, Malengier-Devlies B, Vandermosten L, Pham TT, Mitera T, Possemiers H, Boon L, Opdenakker G, Matthys P, Van den Steen PE: Limitations of neutrophil depletion by anti-Ly6G antibodies in two heterogenic immunological models. Immunol Lett 2019, 212:30–36.

37. Stackowicz J, Jönsson F, Reber LL: Mouse Models and Tools for the in vivo Study of Neutrophils. Front Immunol 2019, 10:3130.

38. Izzy S, Yahya T, Albastaki O, Abou-El-Hassan H, Aronchik M, Cao T, De Oliveira MG, Lu KJ, Moreira TG, da Silva P, et al: Nasal anti-CD3 monoclonal antibody ameliorates traumatic brain injury, enhances microglial phagocytosis and reduces neuroinflammation via IL-10-dependent T(reg)-microglia crosstalk. Nat Neurosci 2025, 28:499–516.

39. Loane DJ, Kumar A, Stoica BA, Cabatbat R, Faden AI: Progressive neurodegeneration after experimental brain trauma: association with chronic microglial activation. J Neuropathol Exp Neurol 2014, 73:14–29.

40. Ertürk A, Mentz S, Stout EE, Hedehus M, Dominguez SL, Neumaier L, Krammer F, Llovera G, Srinivasan K, Hansen DV, et al: Interfering with the Chronic Immune Response Rescues Chronic Degeneration After Traumatic Brain Injury. J Neurosci 2016, 36:9962–9975.

41. Weckbach S, Neher M, Losacco JT, Bolden AL, Kulik L, Flierl MA, Bell SE, Holers VM, Stahel PF: Challenging the role of adaptive immunity in neurotrauma: Rag1(-/-) mice lacking mature B and T cells do not show neuroprotection after closed head injury. J Neurotrauma 2012, 29:1233–1242.

42. Mencl S, Hennig N, Hopp S, Schuhmann MK, Albert-Weissenberger C, Sirén AL, Kleinschnitz C: FTY720 does not protect from traumatic brain injury in mice despite reducing posttraumatic inflammation. J Neuroimmunol 2014, 274:125–131.

43. Zhang L, Ding K, Wang H, Wu Y, Xu J: Traumatic Brain Injury-Induced Neuronal Apoptosis is Reduced Through Modulation of PI3K and Autophagy Pathways in Mouse by FTY720. Cell Mol Neurobiol 2016, 36:131–142.

44. Nemeth DP, Liu X, Chen L, Hawkins MR, Ali HS, Lapid MG, Farsian V, Kim A, Saez J, Maxey G, et al: Temporally-regulated genetic access to IL-1β-expressing cellular networks in homeostasis and following peripheral or central immune stimuli. Brain Behav Immun 2026, 131:106177.

