## Supplemental Figures 1-7 for "Impact of innate immune activation on T cell dynamics and functional recovery following traumatic brain injury"

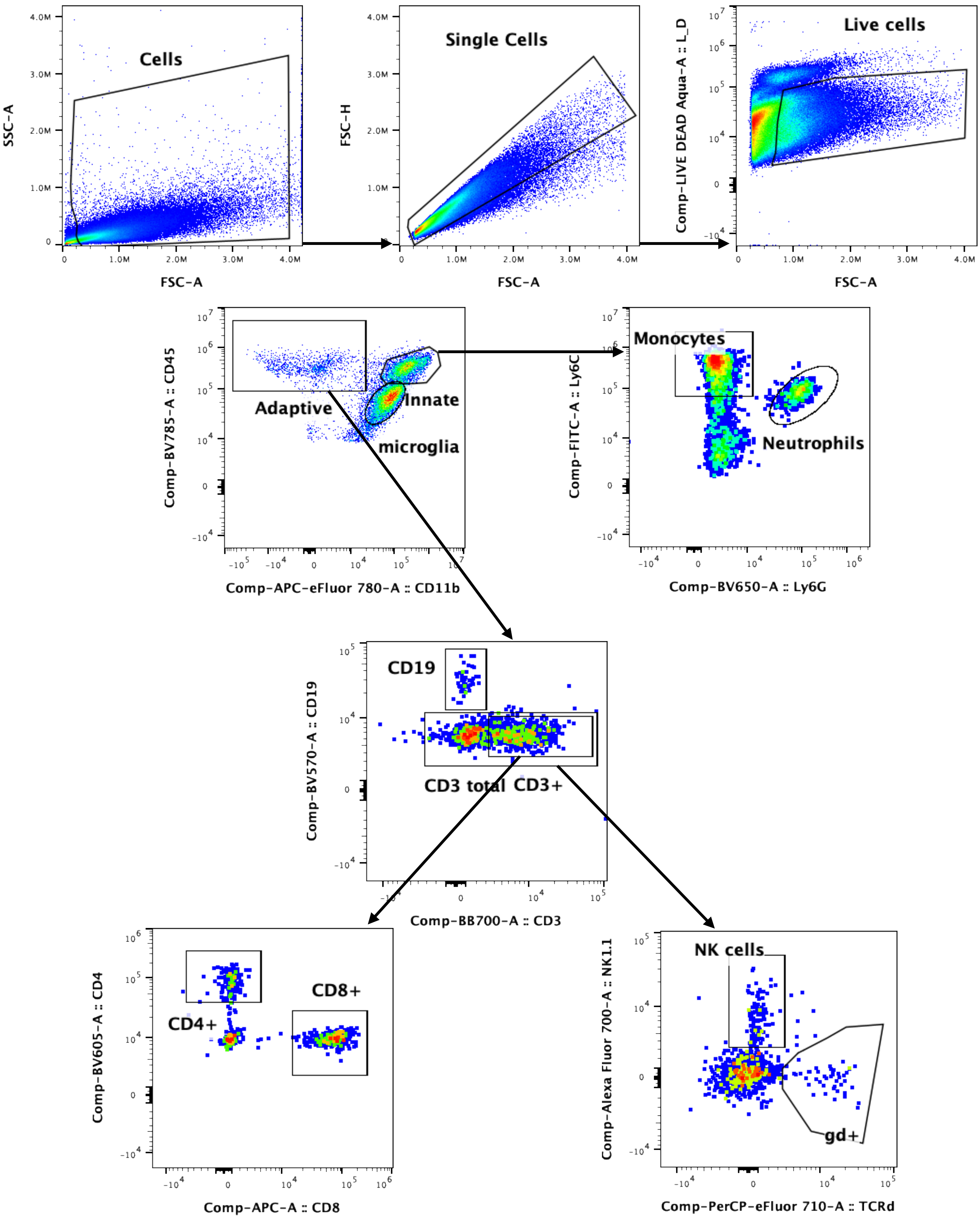


**Supplementary Figure 1. Gating strategy for infiltrating innate and adaptive immune cells in the brain following controlled cortical impact (CCI).**

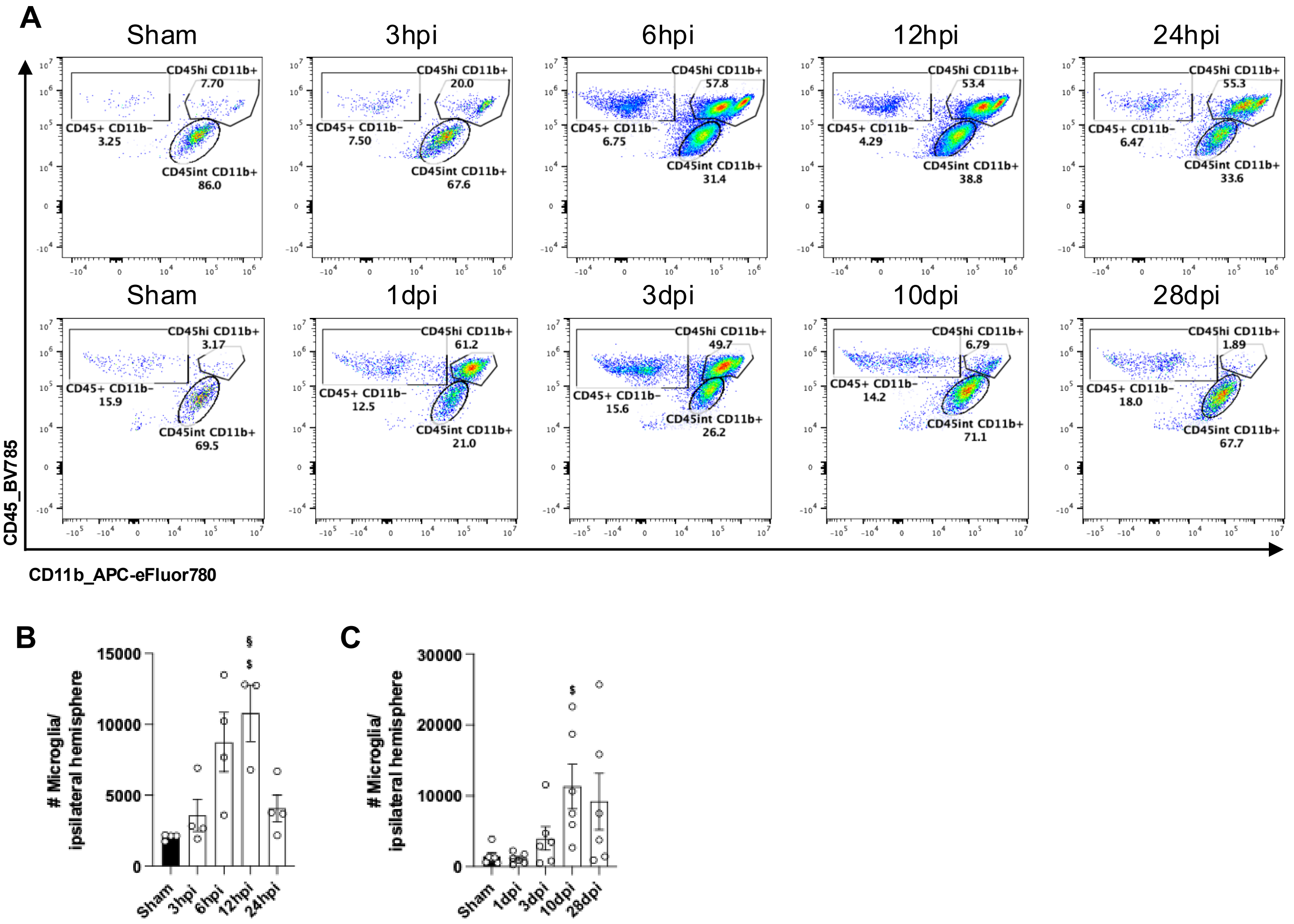


**Supplementary Figure 2. Microglia in the injured brain at acute and chronic time points.** (A) Flow plots used to quantify the microglial population (CD45int CD11b⁺). (B) The bar graphs illustrate the absolute numbers of microglia at the acute time point (3, 6, 12, and 24 hours post injury). Data= Mean ± SEM (n=3-4 per time point); differences vs. sham *p <0.05, **p <0.01, ***p <0.001, differences vs. 3hpi $p <0.05, $$p <0.01, $$$p <0.001, differences vs. 6hpi #p <0.05, ##p <0.01, ###p <0.001, differences vs. 12hpi §p <0.05, §§p <0.01, §§§p <0.001 by one-way ANOVA with Tukey's multiple comparisons test. (C) Microglia at sub-acute and chronic time points (1, 3, 10, 28 days post-injury (dpi)). Data= Mean ± SEM (n=6 per time point); differences vs. sham *p <0.05, **p <0.01, ***p <0.001, differences vs. 1dpi $p <0.05, $$p <0.01, $$$p <0.001, differences vs. 3dpi #p <0.05, ##p <0.01, ###p <0.001, differences vs. 10dpi §p <0.05, §§p <0.01, §§§p <0.001 by one-way ANOVA with Tukey's multiple comparisons.

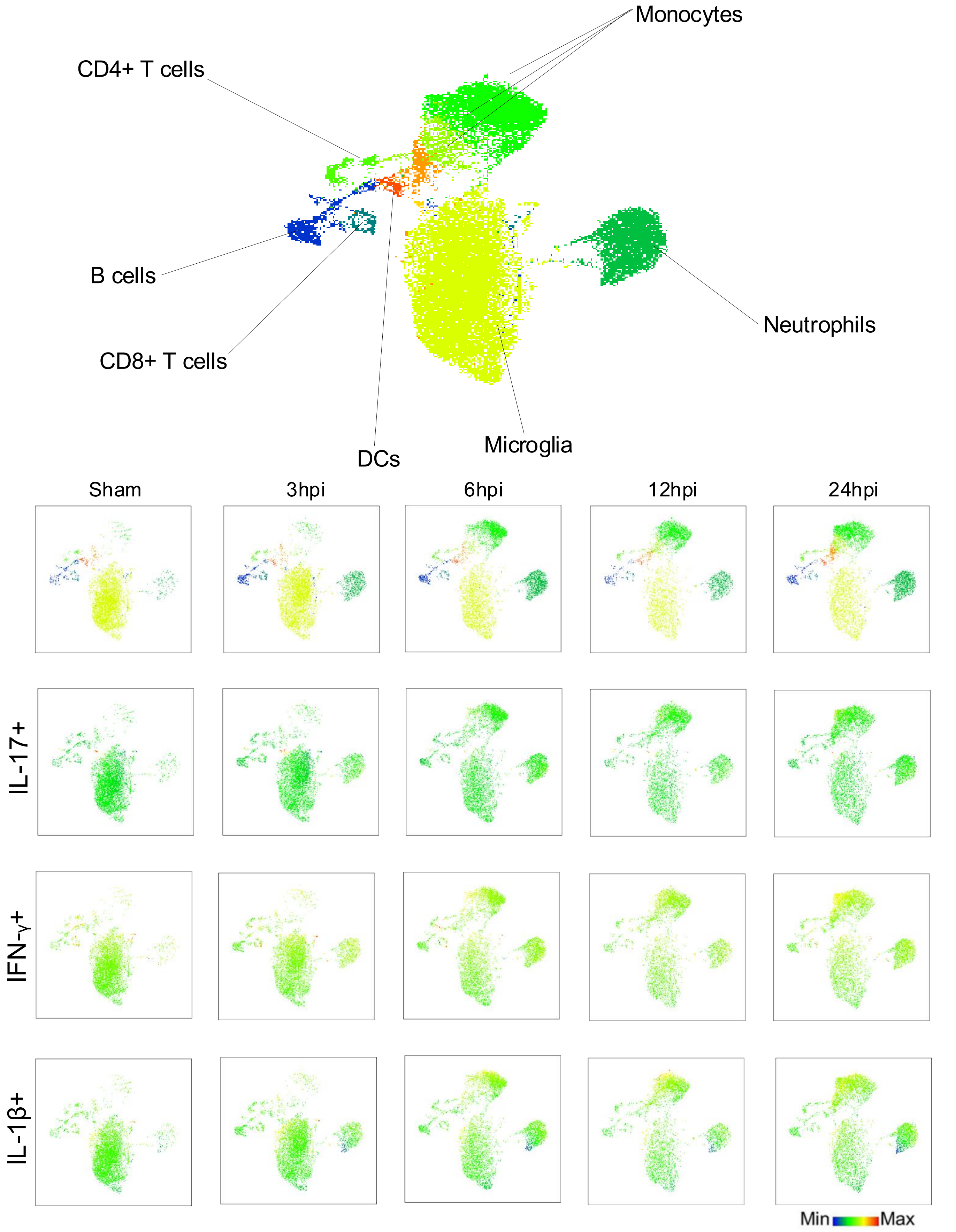


**Supplementary Figure 3. UMAP analysis of innate and adaptive immune cells in the injured brain at acute time points.** (A) UMAP of pooled immune cells from sham and CCI brains across acute time points showing major clusters corresponding to microglia, monocytes, neutrophils, dendritic cells (DCs), $CD4^+$ T cells, $CD8^+$ T cells, and B cells. Split UMAPs show the distribution of cells from Sham, 3, 6, 12, and 24 h post-injury (hpi). Feature plots show IL-17, IFN-γ, and IL-1β signal across the same representation, with colour scale indicating low to high expression.

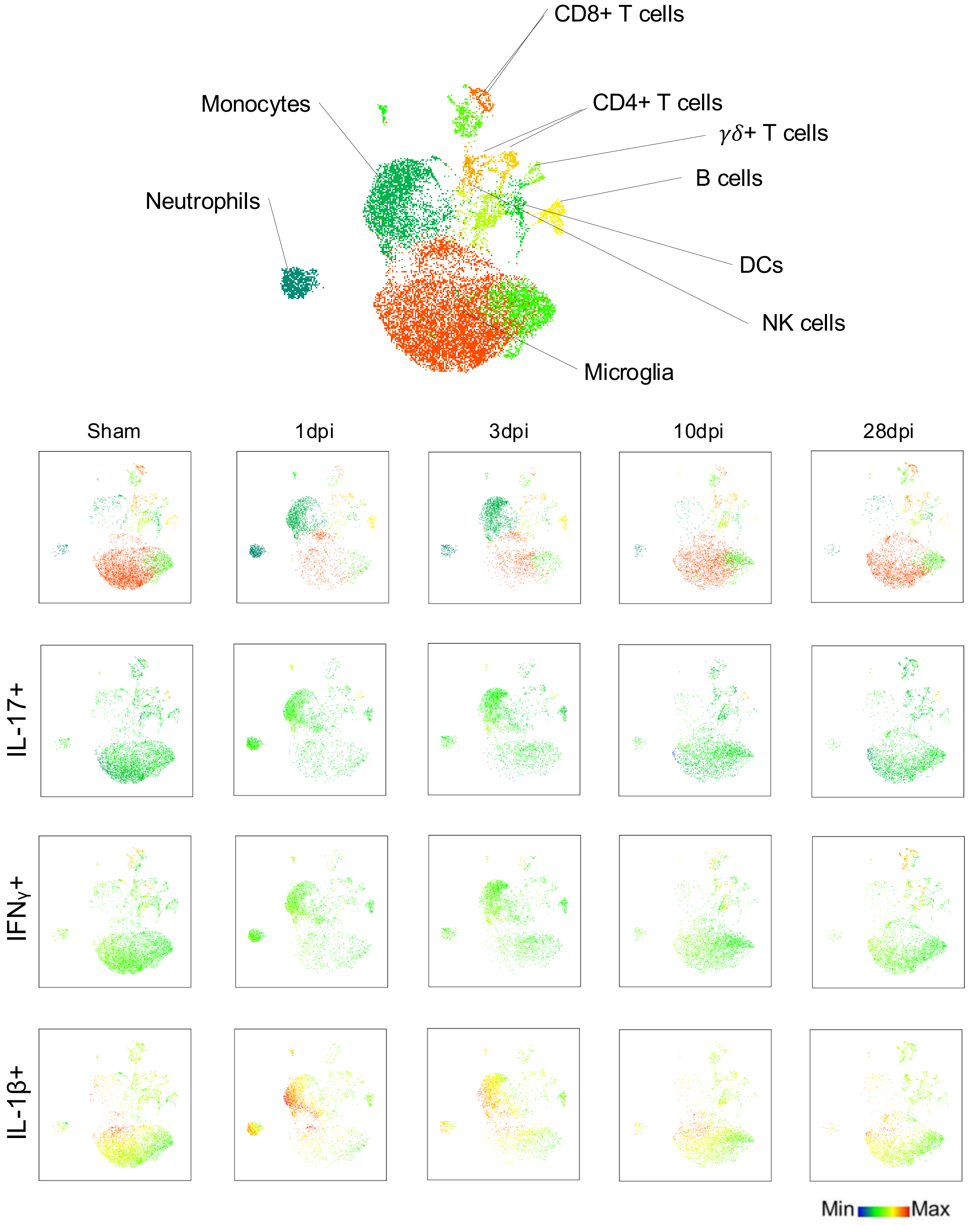


**Supplementary Figure 4. UMAP analysis of innate and adaptive immune cells in the injured brain at sub-acute time and chronic time points.** (A) UMAP of pooled immune cells from sham and CCI brains across acute and sub-acute time points, showing major clusters corresponding to microglia, monocytes, neutrophils, dendritic cells (DCs), natural killer like T (NK-T) cells, B cells, $CD4^+$ T cells, $CD8^+$ T cells, and $\gamma\delta^+$ T cells. Split UMAPs show the distribution of cells from Sham, 1, 3, 10, and 28 dpi. Feature plots show IL-17, IFN-γ, and IL-1β signal across the same representation, with colour scale indicating low to high expression.

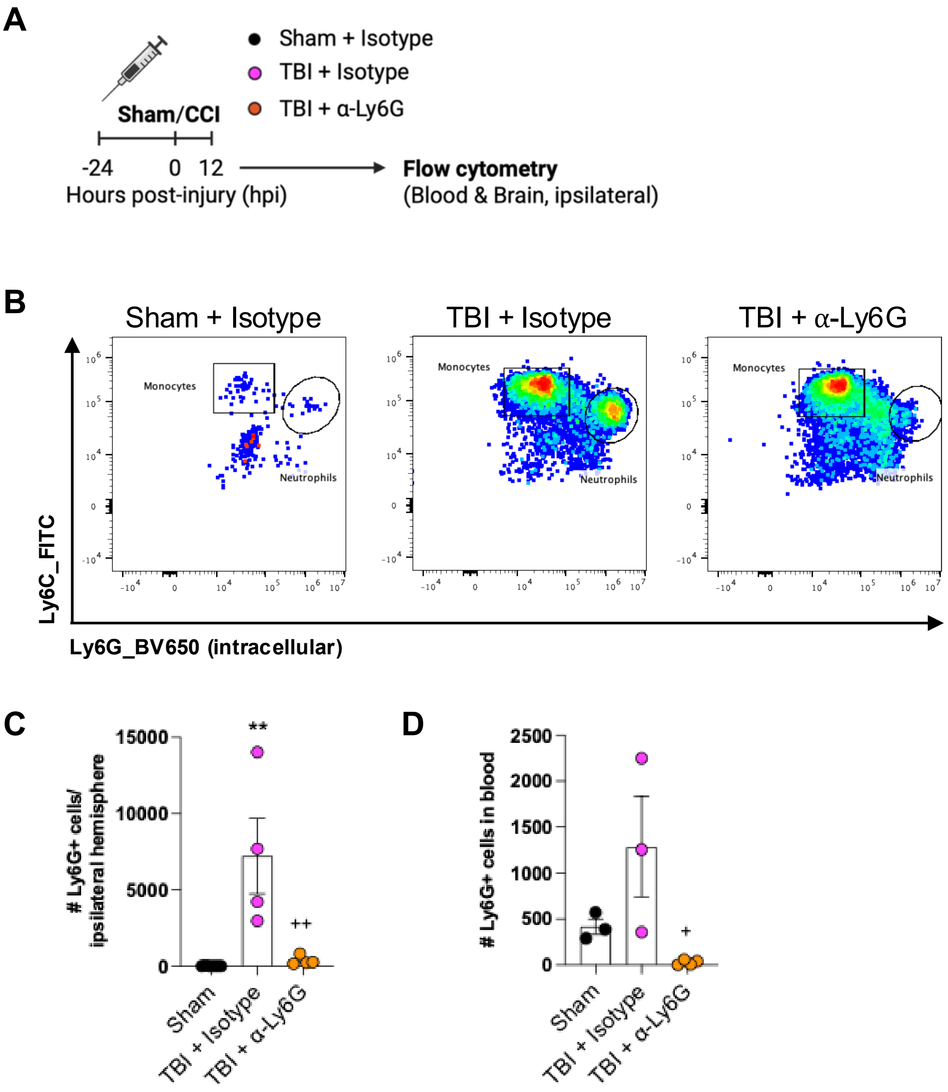


**Supplementary Figure 5. Pilot study for the neutralization of neutrophils.** (A) Experimental design. (B) Representative flow cytometry plots of the group are presented. (C) The bar graph illustrates the absolute number of neutrophils present in the brain and (D) blood. Data = Mean ± SEM (n=3 per group); *p <0.05, **p <0.01, ***p <0.001 by one-way ANOVA, with Tukey's multiple comparisons.

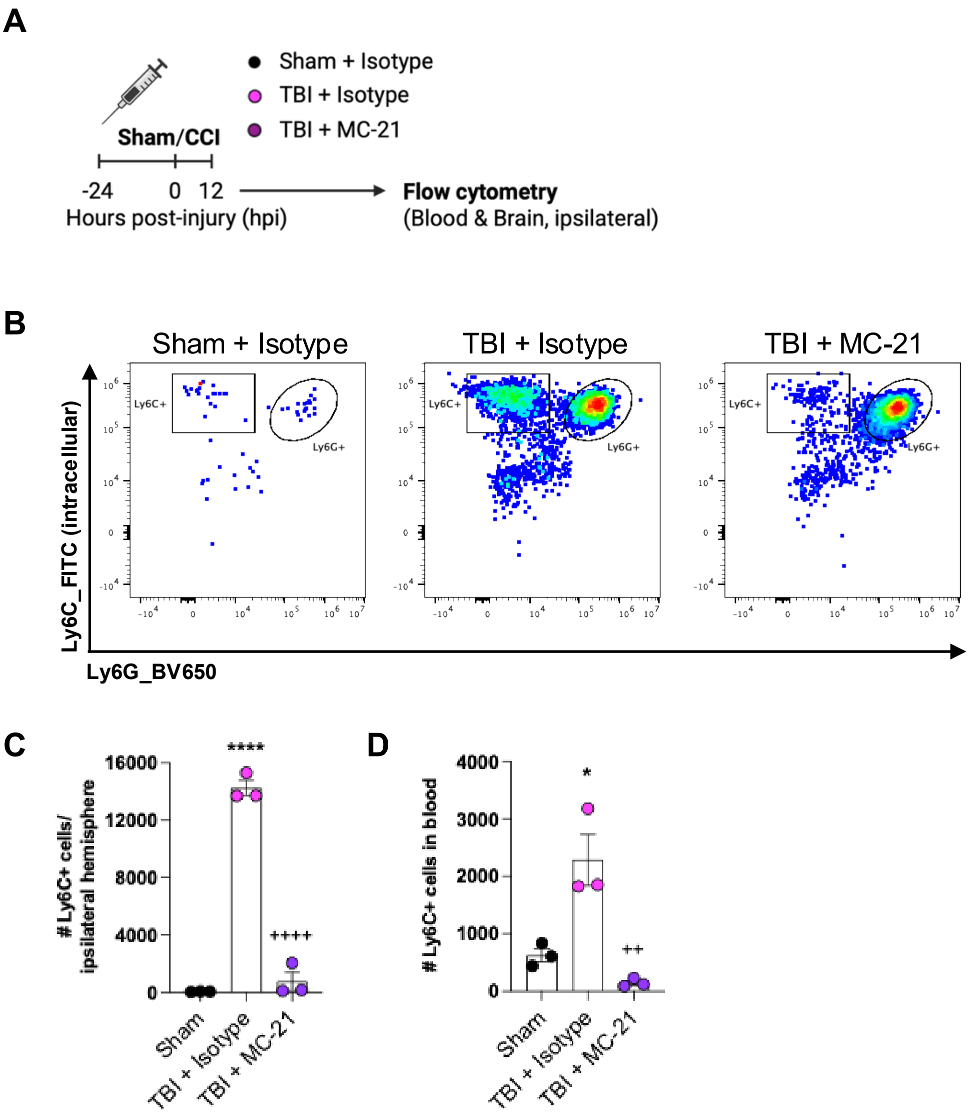


**Supplementary Figure 6. Pilot study for the neutralization of monocytes.** (A) Experimental design. (B) Representative flow cytometry plots of the group are presented. (C) The bar graph illustrates the absolute number of monocytes present in the brain and (D) blood. Data = Mean ± SEM (n=3 per group); *p <0.05, **p <0.01, ***p <0.001 by one-way ANOVA, with Tukey's multiple comparisons.

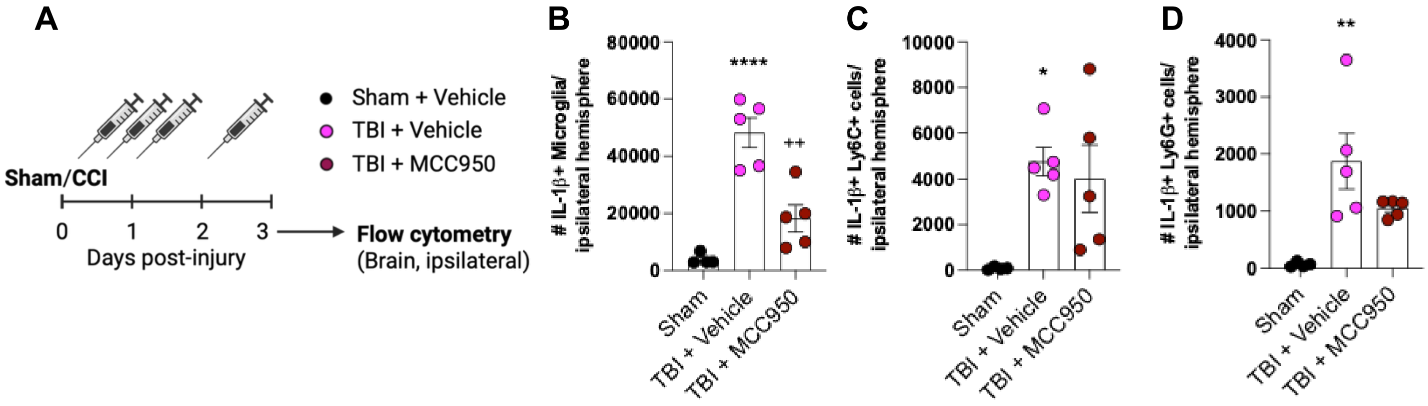


**Supplementary Figure 7. Pilot study for MMC950 treatment.** (A) Experimental design. (B) The bar graph illustrates the absolute number of microglia, (C) Neutrophils, (D) Monocytes. Data = Mean ± SEM (n=3 per group); *p <0.05, **p <0.01, ***p <0.001 by one-way ANOVA, with Tukey's multiple comparisons.
